# Bacteriophages target membrane-anchored glycopolymers to promote host cell lysis and progeny release

**DOI:** 10.1101/2025.06.24.661397

**Authors:** Amelia C. McKitterick, Evan W. Lyerly, Thomas G. Bernhardt

## Abstract

Most bacteriophages lyse their host cell to release progeny virions. Double-stranded DNA phages typically promote host lysis using a holin-endolysin system. Holins form pores in the cytoplasmic membrane allowing endolysins access to the peptidoglycan (PG) cell wall, which they degrade to weaken the cell envelope and promote osmotic lysis. Phages that infect Proteobacteria also encode a spanin in their lysis cassette that functions to disrupt the host outer membrane. The spanin-requirement for cell lysis provided the first clue that the proteobacterial outer membrane confers mechanical rigidity to the cell envelope. Corynebacteria and mycobacteria also build an outer membrane, but it is made of mycolic acids instead of lipopolysaccharides. Here, we investigated whether the mycomembrane presents a mechanical barrier to phage-induced lysis of corynebacteria. In addition to annotated holin and endolysin genes, mycobacteriophages and corynephages were found to encode a membrane protein we call LysZ in their lysis cassettes. Deletion of *lysZ* in the corynephage Cog blocked lysis of its host *Corynebacterium glutamicum*. Surprisingly, disruption of the host mycomembrane did not correct this phenotype. Instead, a genetic analysis revealed that blocking synthesis of membrane-anchored glycopolymers called lipomannans/lipoarabinomannans (LM/LAMs) can restore plaque formation when LysZ is inactivated. This genetic system also identified the likely flippase that transports decaprenyl-linked mannose units to the extracellular side of the membrane for polymerization into LM/LAM. Overall, our results indicate that lipoglycans like LM/LAMs play roles in mechanically stabilizing bacterial envelopes and that phages use LysZ-like factors to overcome this barrier to promote lysis.

**SIGNIFICANCE:** Bacteriophages must break down the cell envelope of their host to induce cell lysis for progeny release. Pathogens like *Mycobacterium tuberculosis* and related corynebacteria build complex cell envelopes that are likely to be especially challenging for phages to disrupt. Here, we identify a novel lysis gene called *lysZ* encoded by many mycobacteriophages and corynephages. Using the model organism *Corynebacterium glutamicum*, we show that LysZ is required to induce host lysis. This requirement is overcome by inactivating genes needed for the synthesis of lipid-anchored glycans called lipomannans/lipoarabinomannans (LM/LAMs). These findings reveal that LM/LAMs are likely to play a mechanical role in stabilizing the cell envelope. LM/LAM biogenesis is therefore an attractive target for drugs that disrupt the integrity of mycobacterial envelopes.

## INTRODUCTION

To release progeny virions at the end of a replication cycle, most bacteriophages induce lysis of their host cell. This abrupt demise of the host bacterium is promoted by phage proteins that disrupt the stress-bearing capabilities of the cell envelope. Internal osmotic pressure then promotes the explosive rupture of the compromised cell. Double-stranded DNA phages typically use a holin-endolysin system to promote cell lysis^1^. During phage replication, holin proteins accumulate in the cytoplasmic membrane^2^. Depending on the phage, the endolysin either accumulates in the cytoplasm or in an inactive conformation in the membrane. At a genetically programmed time in the replication cycle optimized for progeny dispersal, the holins form pores in the membrane to release the endolysins from the cytoplasm or membrane, allowing them to reach their substrate, the peptidoglycan (PG) cell wall. The released endolysins then degrade the wall to trigger osmotic lysis.

In the case of phages that infect diderm Proteobacteria like *Escherichia coli*, the holin and endolysin alone are not sufficient to promote efficient lysis. A third factor called a spanin is needed to destabilize the outer membrane of these bacteria (**Fig. 1A**). Spanins are often encoded just downstream of the holin and endolysin genes in a phage genomic locus referred to as the lysis cassette^3^. In the model bacteriophage λ, the spanin is formed by two proteins Rz and Rz1, with Rz1 encoded entirely within the Rz gene in a different reading frame. Rz is an inner membrane protein and Rz1 is an outer membrane lipoprotein. When the PG layer is destroyed by the endolysin, the Rz and Rz1 proteins are thought to form a transenvelope complex that connects the inner and outer membrane and fuses them to provide the final disruptive blow to the cell envelope^4,5^. Under conditions where the outer membrane is stabilized, such as in the presence of Mg^2+^, or in the absence of significant sheer forces, mutant phage lacking a spanin fail to complete lysis^6^. The host cell loses its cell shape as expected when the endolysin degrades the PG layer, but the spheroplast is stabilized by the intact outer membrane, which prevents progeny release. This spanin requirement for phage-induced cell lysis provided the first indication that the outer membrane of Gram-negative bacteria provides mechanical stability to the cell envelope in addition to its well-known role as a permeability barrier that provides these organisms with their high intrinsic resistance to antibiotics^7^.

**Figure 1.**
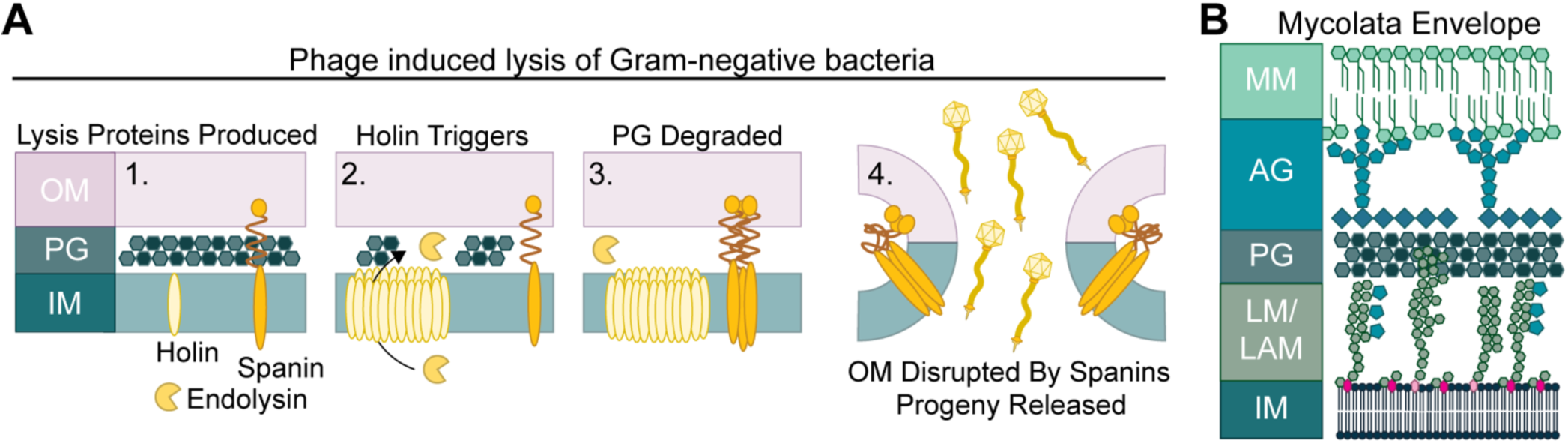
Gram-negative cell lysis and the mycolata envelope. **A**. Cartoon detailing the steps of phage-induced lysis of a Gram-negative cell. (Step 1) Production of the phage lysis proteins: holin, endolysin, and spanin. (2) Holin triggering leads to formation of large pores in the membrane that allow the endolysin to enter the periplasmic space. (3) The endolysin degrades the peptidoglycan (PG) cell wall, allowing spanin multimerization. (4) Multimerized spanins induce fusion of the inner membrane (IM) and outer membrane (OM), leading to lysis and phage progeny release. **B**. Diagram of the mycolata envelope that surrounds bacteria of the Mycobacteriales order. IM, inner membrane; LM/LAM, lipomannan and lipoarabinomannan; PG, peptidoglycan; AG, arabinogalactan; MM, mycomembrane; OM, outer membrane.

Like the Proteobacteria, bacteria belonging to the order Mycobacteriales such as the pathogen *Mycobacterium tuberculosis* and the related model organism *Corynebacterium glutamicum* (*Cglu*) are diderm. They possess a second membrane that covers a complex, multi-layered cell envelope called the mycolata envelope consisting of a PG layer decorated with another polysaccharide called arabinogalactan (AG)^8^ (**Fig. 1B**). However, instead of the phospholipids and lipopolysaccharides that make up the Gram-negative outer membrane, this membrane is made of lipid species called mycolic acids. We were therefore curious whether phages that infect members of the Mycobacteriales encode spanin-like factors to overcome this unique mycomembrane barrier. Indeed, some mycobacteriophages encode an esterase called LysB that cleaves linkages between mycolic acids and the AG layer to disrupt the mycomembrane and promote progeny release^9,10^. In this report, we investigate whether corynephages also use mycomembrane disruptors to induce host lysis.

Genome analysis revealed that instead of *lysB* genes, phages that infect *Cglu* encode a membrane protein of unknown function that we call LysZ in their lysis cassette. LysZ is widespread among not only corynephages but also phages that infect a variety of bacteria within the Actinomycetota phylum to which the Mycobacteriales order belongs. Inactivation of LysZ in a model corynephage called Cog that infects *Cglu* was found to block cell lysis. Surprisingly, disruption of the mycomembrane did not rescue the plaque formation defect of a Δ*lysZ* phage. Rather, a genetic analysis revealed that blocking the synthesis of lipid-linked polysaccharides called lipomannans (LMs) and lipoarabinomannans (LAMs) restored plaque forming ability to phages inactivated for LysZ. Notably, this analysis identified the likely flippase that transports decaprenyl-phosphomannose precursors across the membrane for LM/LAM biogenesis. Additionally, mutants defective for late stages of LM/LAM synthesis were found to have growth and shape defects that are likely to be caused by depletion of the decaprenyl lipid carrier. By analogy with the spanin results, we infer that like the outer membrane of Gram-negative bacteria, LM/LAMs play a role in providing mechanical stability to the cell envelopes of the bacteria that produce them.

## RESULTS

### Genome analysis identifies lysZ as a potential lysis gene in the Corynebacteriales

To understand how corynephages dismantle the envelope of *Cglu*, the genomes of the corynephages CL31 and Cog were analyzed to identify lysis genes. Like most phages, CL31 and Cog have a modular genome organization in which genes with related functions tend to be encoded in the same regions of the genome. Proximal to the tail genes, CL31 and Cog encode what appears to be a three-gene lysis cassette composed of *lysA*, *hol*, and a gene of unknown function (**Fig. 2A**). The *lysA* gene encodes a predicted N-acetylmuramoyl-L-alanine amidase, an enzymatic activity for PG cell wall degradation, indicating that LysA is likely to be the phage endolysin. The *hol* gene is predicted to encode a holin of the Phage_r1t family, members of which contain two transmembrane (TM) domains separated by a short β-turn^11,12^. The last gene in the lysis cassette encodes a protein of unknown function that contains a predicted disordered N-terminal domain, a TM helix, a long α-helical body, a flexible, proline-rich linker, and a C-terminal helix (**Fig. 2A and B**). Based on the results presented below, we have called this protein LysZ. Sequence similarity searches in NCBI identify LysZ sequences in corynephage phi16 and a putative temperate phage in *Cglu* ATCC 13870. Using proximity to endolysin-encoding genes and the predicted structural features of LysZ, we identified *lysZ*-like genes in diverse phages that infect a range of bacteria in the Actinomycetota phylum (**Fig. 2A and B**). These phages include those that infect members if the Mycobacteriales order such as *Mycobacterium smegmatis* and *Gordonia terriae*, as well as bacteria in other related orders such as the Micrococcales. In many of these phages, the *lysZ*-like gene is encoded downstream of a gene predicted to encode a protein with four TM domains (**Fig. 2A**). We have named these proteins LysY due to their location in the putative lysis cassettes of many phages. In this report, we focused on determining the function of the LysZ proteins.

**Figure 2.**
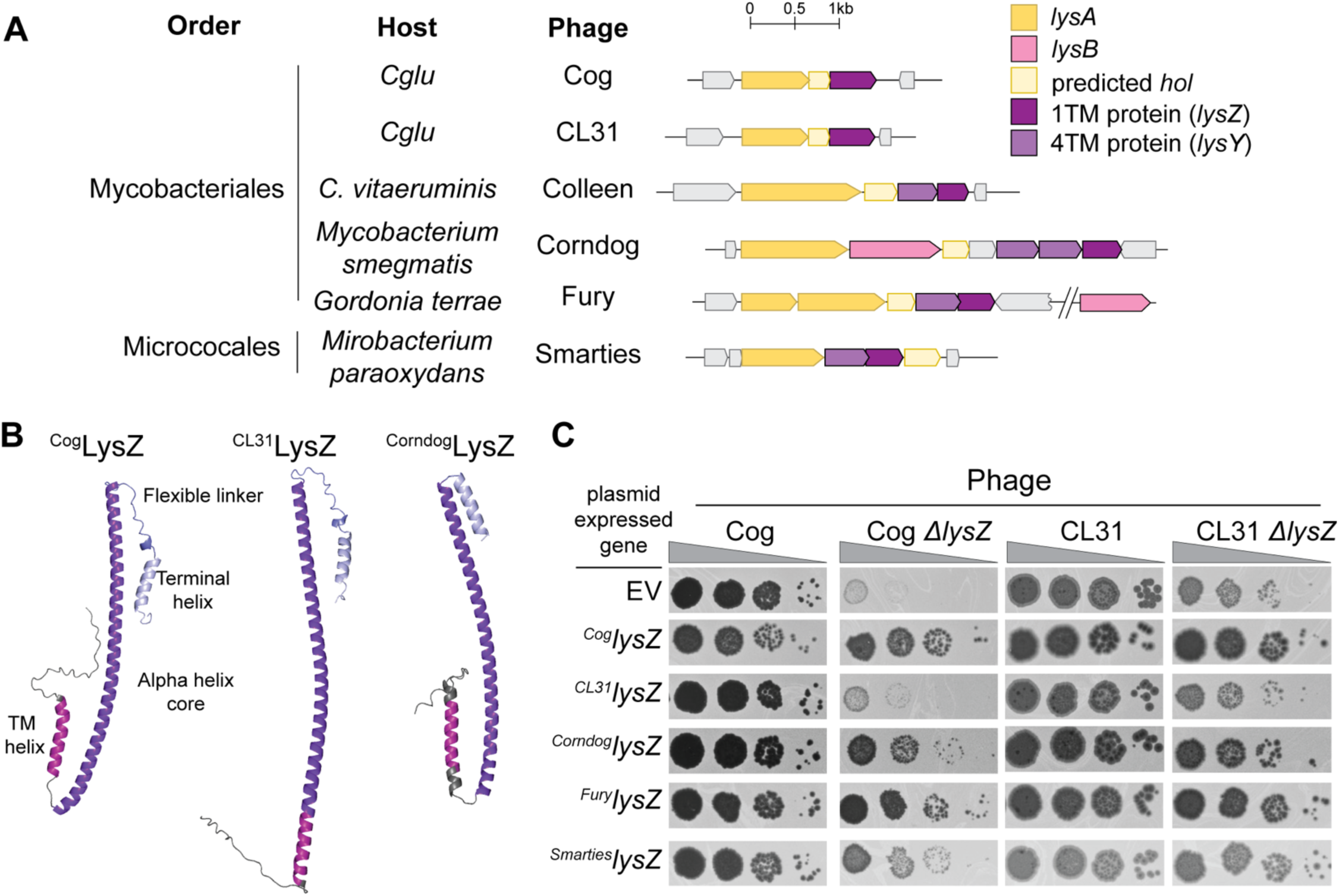
LysZ is required for efficient corynephage plaque formation. **A**. Diagrams of lysis cassettes from the corynephage Cog and other actinophages. The genes encoded in the cassettes are: *lysA* endolysin, *lysB* esterase, *hol* holin, and predicted lysis genes *lysZ* and *lysY*. **B**. Alphafold2 models of LysZ structures from the indicated phages with structural features highlighted. **C**. Tenfold serial dilutions of the indicated phages were spotted on a lawn of wild-type *Cglu* harboring the indicated gene on a multicopy plasmid under control of an IPTG-inducible *tac* promoter (P*tac*) and a modified *riboE1* riboswitch (*riboE1\**) that allows translation in the presence of the inducer theophylline. Expression was induced by the addition of IPTG (100 µM) and theophylline (1 mM) to the soft agar. EV denotes empty vector.

### Cog and CL31 phages require LysZ for normal plaque formation

To determine if *lysZ* is required for host lysis, mutant CL31 and Cog phage deleted for *lysZ* were constructed using CRISPR-Cas engineering. CL31 Δ*lysZ* phage formed smaller plaques than wild-type CL31 on a lawn of the wild-type *Cglu* strain MB001 whereas a Cog mutant lacking *lysZ* had a severe plating defect (**Fig. 2C, S1A-B**). The plating defect of Δ*lysZ* Cog phage was complemented by the expression of the cognate *lysZ* gene from a plasmid. Additionally, ^Cog^*lysZ* complemented the small plaque phenotype of the CL31 Δ*lysZ* phage. A construct expressing ^CL31^*lysZ* failed to restore the plaquing ability of CL31 Δ*lysZ* or Cog Δ*lysZ* for unknown reasons. However, *lysZ* alleles from the distantly related phages Corndog, Fury, and Smarties that infect *M. smegmatis G. terrae*, and *Mirobacterium paraoxydans*, respectively were able to complement the plating defects of the *lysZ* deletion mutants of Cog and CL31 (**Fig. 2C, S1A-B)** The mutant phenotypes and complementation results indicate that LysZ plays an important and likely conserved role in the propagation of phages that infect members of the Mycobacteriales and their relatives.

### LysZ is required for host cell lysis but does not target the mycomembrane

In addition to their defects in plaque formation, Cog and CL31 phages deleted for *lysZ* displayed a defect in inducing cell lysis in a liquid infection assay (**Fig. 3A, Fig. S2**). As with the plaquing phenotypes, the lysis defect in liquid medium was complemented by inducing the expression of *lysZ* alleles from a plasmid at the time of infection. These lysis defects were associated with a decrease in phages released during infection (**Fig. 3B**). To further investigate the defect caused by *lysZ* inactivation, we used time-lapse microscopy to follow the changes in cell morphology and cell lysis induced by wild-type Cog or the Δ*lysZ* mutant following the infection of cells with a high multiplicity of infection (MOI). Cells began lysis at about 90 minutes post infection with wild-type Cog, with the majority of cells being lysed after 120 minutes (**Fig. 3C, Movie 1**). By contrast, lysis was not observed up to five hours after infection with the Cog Δ*lysZ* phage. Instead, the cells were observed to progressively become more bulbous and inflated at the poles (**Fig. 3C, Movies 2 and 3**), suggesting that the cell envelope was being partially deconstructed by the phage to cause the loss in cell shape, but that the final step of osmotic lysis was blocked. This result is reminiscent of the observation of stable spheroplasts of *E. coli* formed following infection with phage λ inactivated for its spanin^6^. To determine if the mycomembrane is stabilizing the envelope of *Cglu* cells infected with the Cog Δ*lysZ* phage, we tested whether the loss of LysZ function could be complemented by production of a mycobacteriophage LysB esterase that cleaves mycolic acid-AG linkages^10^. Surprisingly, LysB production did not restore the ability of the Cog Δ*lysZ* phage to form plaques (**Fig. 3D**). Unlike mycobacteria, mutants of *Cglu* lacking the mycomembrane are viable^13^. We therefore also tested whether the Cog Δ*lysZ* phage could form plaques on lawns of *Cglu* mutants inactivated for the *afhA* or *ostA* genes needed for mycolic acid biogenesis^14^. Again, to our surprise, preventing mycomembrane assembly did not rescue the plaque formation defect of the Cog Δ*lysZ* phage (**Fig. 3E**). A similar test for CL31 Δ*lysZ* phage could not be performed because our prior results indicate that CL31 requires the mycomembrane for infection whereas Cog does not^14^. Based on these results, we conclude that LysZ is required by corynephages to promote efficient host cell lysis and that its function is unrelated to mycomembrane disruption.

**Figure 3.**
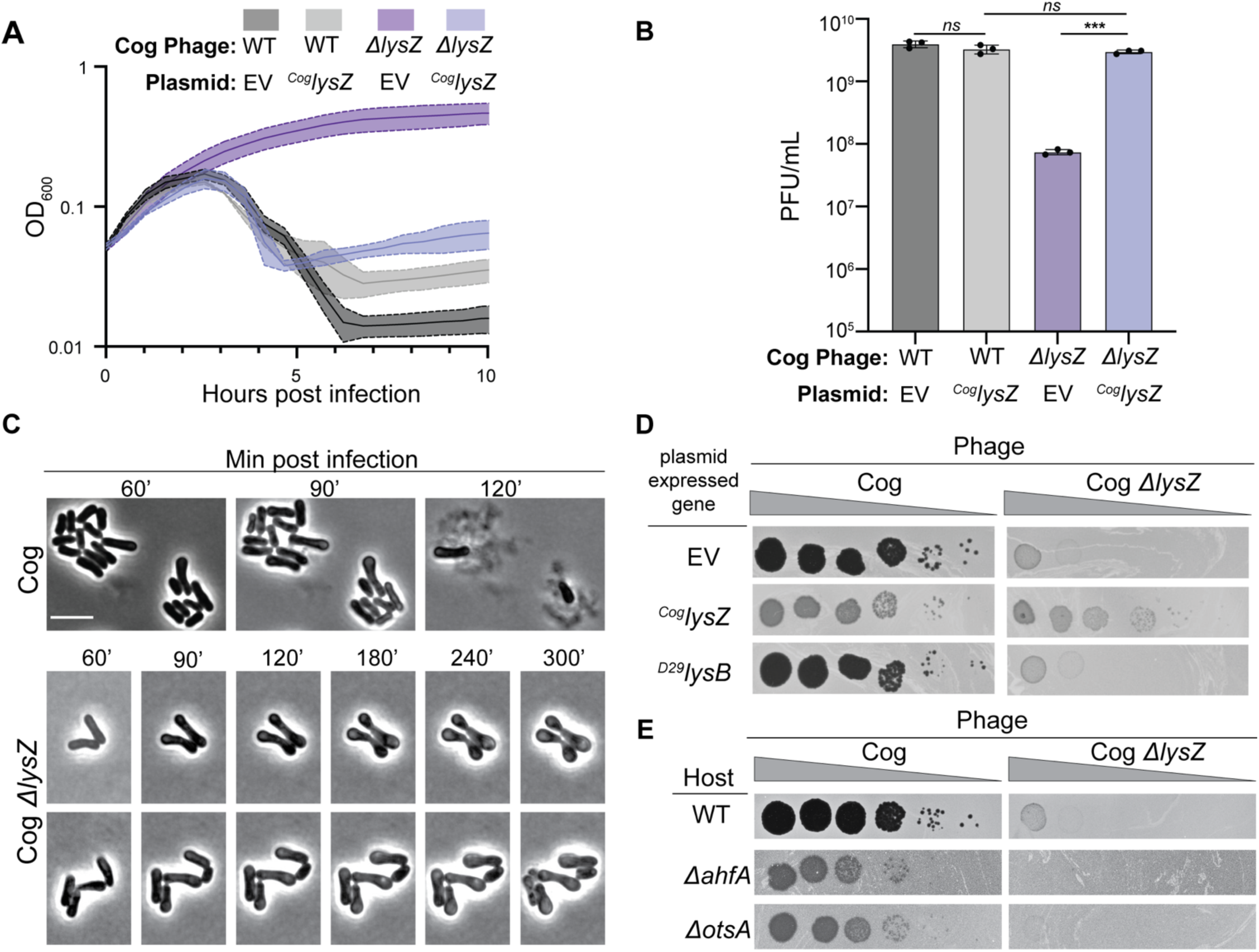
LysZ is needed to induce host cell lysis. **A**. Cells of *Cglu* harboring the indicated expression plasmid were infected with the indicated phage at an MOI of 2.5. Expression of the indicated genes was induced with IPTG (100 µM) and theophylline (1 mM) at the time of infection. Growth of the infected culture was followed by monitoring OD600. Shaded regions indicate standard deviation, and solid lines indicate the mean from three independent experiments. **B.** Enumeration of phages released 120 minutes post infection of the listed strains with the listed phages. Expression of the indicated genes was induced with IPTG (100 µM) and theophylline (1 mM) at the time of infection. Dots represent individual data points, bar heights represent the mean, and error bars indicate standard deviation of three independent experiments. Significance was determined by a two-tailed t test. *ns,* not significant; ***p<0.0001. **C**. Time-lapse microscopy of *Cglu* infected at an MOI of 10 with the indicated Cog variant and immobilized on agarose pads. Images of *Cglu* infected with Cog *ΔlysZ* show two representative sets of cells. Bar equals 5 µm. **D.** Tenfold serial dilutions of the indicated phages were spotted on a lawn of wild-type *Cglu* harboring the indicated gene on a multicopy plasmid induced as in Fig. 2. **E.** Tenfold serial dilutions of the indicated phages were spotted on lawns of the indicated *Cglu* hosts.

### Mutants defective for decaprenyl-phosphomannose synthesis rescue the Cog *Δ*lysZ plating defect

To identify the layer of the cell envelope targeted by LysZ, we took advantage of the finding that overexpression of ^Cog^*lysZ* is toxic (**Fig. 4A-C**). Reasoning that spontaneous suppressors resistant to ^Cog^LysZ production might identify mutants that alter or eliminate the envelope layer that LysZ is required to disrupt, we isolated survivors following the induction of ^Cog^*lysZ* expression from a plasmid. Among the survivors of ^Cog^*lysZ* expression, we anticipated a high frequency of isolates harboring plasmids defective for ^Cog^*lysZ* expression in addition to the mutants of interest. The survivors were therefore screened using two additional assays. The first screen was for isolates hypersensitive to the detergent sodium dodecyl sulfate (SDS). Here, we expected isolates with defective plasmids to retain normal SDS-resistance. However, we expected that mutants with an altered cell envelope conferring LysZ resistance might also have a compromised permeability barrier rendering them sensitive to SDS. We therefore patched survivors of ^Cog^*lysZ* expression on media with or without SDS and selected those that displayed SDS hypersensitivity. These isolates were then tested for their ability to serve as permissive hosts for the Cog Δ*lysZ* phage. Finally, to map the mutations responsible for the desired phenotypes, mutant isolates that allowed plaque formation by the Cog Δ*lysZ* phage were analyzed by whole genome sequencing.

**Figure 4.**
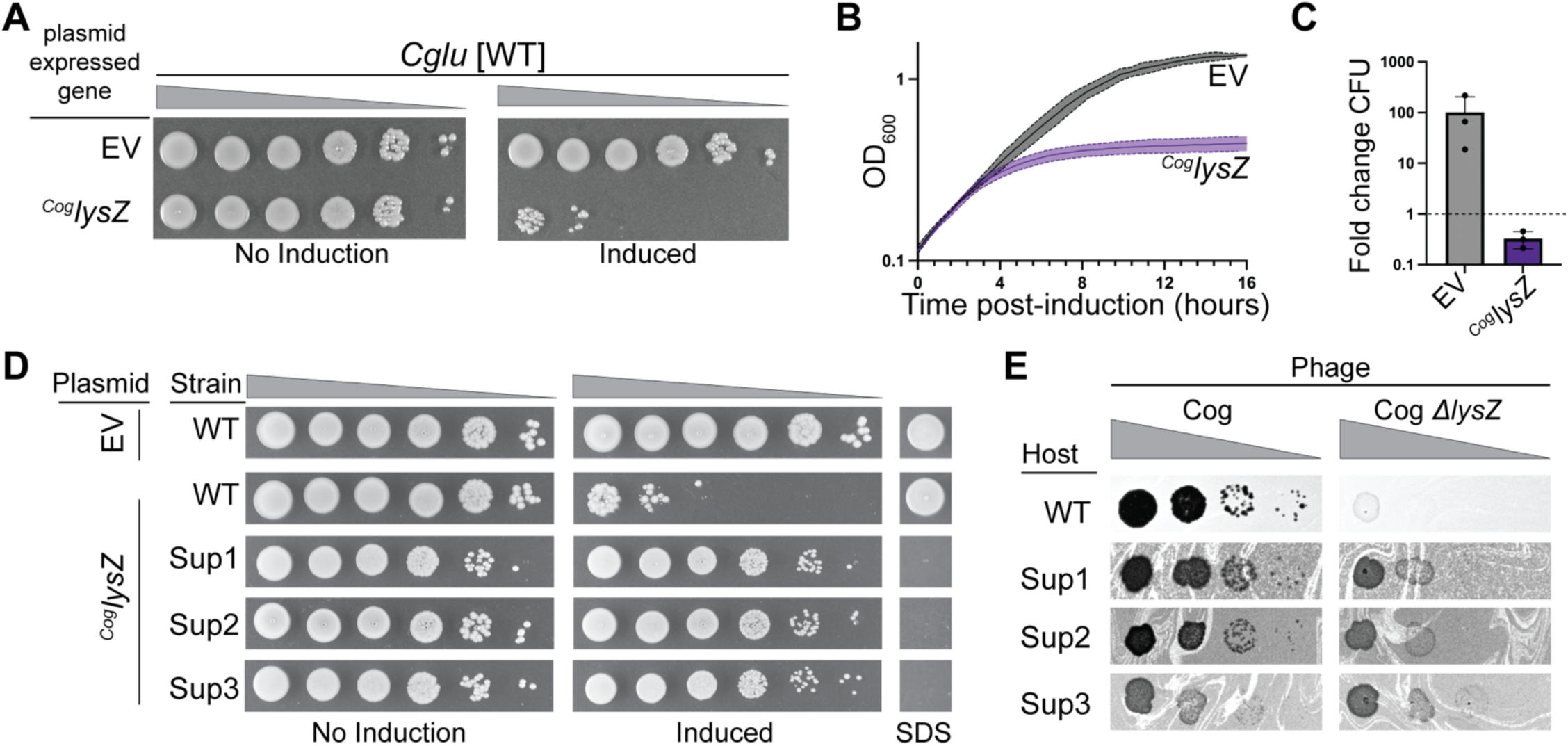
^Cog^LysZ overexpression is lethal. **A**. Wild-type *Cglu* with an empty vector (EV) or ^Cog^*lysZ* containing expression plasmid were serially diluted and plated on agar with and without IPTG (100 µM) and theophylline (1 mM) for induction of gene expression from the plasmid. **B**. Growth of the same strains in (**A**) was followed by monitoring culture OD600 after induction with IPTG (100 µM) and theophylline (1 mM). Shaded regions indicate standard deviation, and solid lines indicate the mean from three independent experiments. **C.** Change in colony forming units (CFU) of the indicated strains after overnight induction as in (**C**). Dots represent individual data points, bar heights represent the mean, and error bars indicate standard deviation of three independent experiments. Significance was determined by a two-tailed t test. **D.** Tenfold dilutions of the indicated *Cglu* expression strain spotted with or without inducers as in (**A**). To the right, SDS sensitivity was monitored. Cultures normalized to OD600 = 1 were spotted on agar containing 0.0075% SDS. Sup1 = *cgp_0855*::*IS*, Sup2 = *manB*, and Sup3 = *ppmC*. **E.** Tenfold serial dilutions of the indicated phages were spotted on lawns of the indicated *Cglu* hosts.

Three mutant isolates were identified through this screen (**Fig. 4D-E**). One of the suppressor mutations that promoted LysZ resistance and restored plaque forming ability to the Cog Δ*lysZ* phage was an insertion element that interrupted the *cgp_0855* gene (**Table S1**). This gene is upstream of *manA*, which encodes an enzyme that interconverts fructose-6-phosphate into mannose-6-phosphate for ultimate use in the synthesis of glycolipids like phosphatidylinositol mannosides (PIMs) and LM/LAM glycopolymers in the cell envelope^15^ (**Fig. 5A**). The second suppressor harbored three mutations, with one being a single nucleotide change in *manB* (also referred to as *pmmA*), which encodes the enzyme acting after ManA in the synthesis pathway for generating GDP-mannose for use in envelope biogenesis^16^ (**Fig. 5A**). Finally, the third suppressor had a mutation mapping to *ppmC*, which encodes the enzyme that uses GDP-mannose created by ManC as a substrate to generate the lipid-linked sugar decaprenyl-phosphomannose (DPM)^17–19^ (**Fig. 5A**). Thus, all the isolates from the selection and screening combination had mutations predicted to disrupt DPM biogenesis.

**Figure 5.**
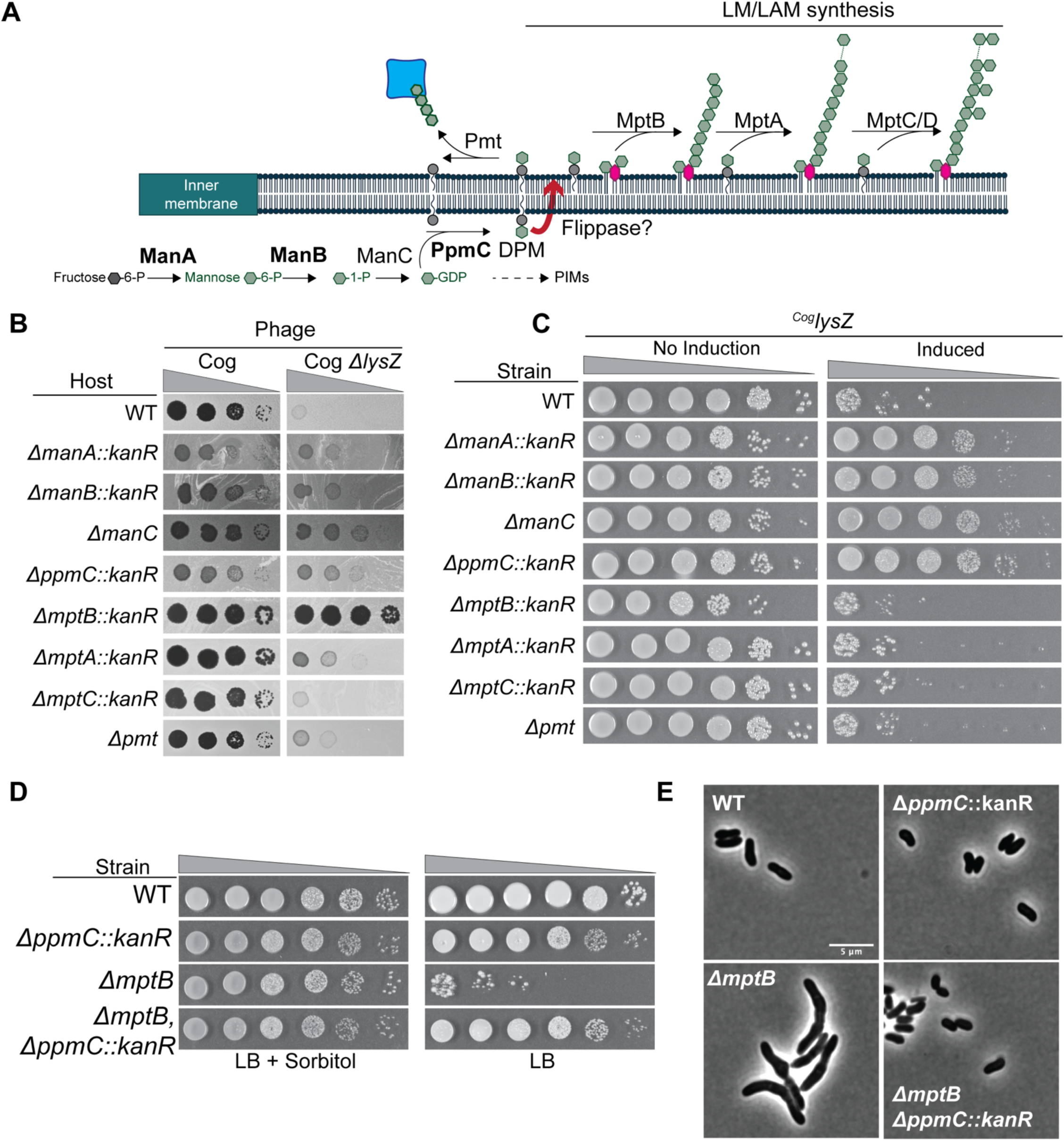
LysZ is dispensable host lysis in strains lacking LM/LAM. **A**. Diagram of the LM/LAM biogenesis pathway. See text for details. Enzymes highlighted in bold indicate steps in the pathway affected by suppressor mutations that alleviate ^Cog^*lysZ* toxicity. **B**. Tenfold dilutions of the indicated phage variants spotted on lawns of the indicated mutant *Cglu* hosts. **C.** Tenfold dilutions of the indicated strains harboring the ^Cog^*lysZ* expression plasmid were plated with and without inducer as in Fig. 4A. **D**. Tenfold dilutions of the indicated *Cglu* strains spotted on LB agar with or without sorbitol (9.1%) as indicated. **E**. Phase contrast images of the indicated strains. Bar equals 5 µm.

### Evidence that LysZ is required to overcome an LM/LAM barrier for Cog to elicit lysis

To validate the results from the suppressor selection/screen, we generated *Cglu* deletion mutants inactivated for ManA, ManB, ManC, and PpmC. As expected, each isogenic deletion mutant conferred resistance to ^Cog^*lysZ* expression and rescued the plating defect of the Cog Δ*lysZ* phage (**Fig. 5B, 5C, and S3C**). Inactivation of *cgp_0855* did not allow plaque formation by the Cog Δ*lysZ* phage, suggesting that the phenotype of the original *cgp_0855* mutant isolate resulted from a polar effect of the insertion element on the expression of the neighboring *manA* gene (**Fig. S3A**).

To identify the DPM-related envelope component responsible for inhibiting plaque formation by the Cog Δ*lysZ* phage, we constructed several mutants defective in the conversion of DPM to surface glycans. Following its synthesis, the phosphomannose headgroup of DMP must be externalized by a yet to be identified lipid flippase for its polymerization into mannose-based glycans used for envelope biogenesis. Once flipped, DPM is used to polymerize LM/LAMs on the lipid anchor called acylated phosphatidylinositol mannoside (AcPIM2) or to form a lipomannan variant called LM-B that is linked to the anchor mannosyl-glucuronic acid diacylglycerol (Gl-X)^20^. In both cases, the glycosyltransferase MptB initiates mannan polymerization and the glycosyltransferases MptA and MptC extend the glycan and introduce branches, respectively^21–23^ (**Fig. 5A**). Deletion of *mptB* robustly restored plaque formation by Cog Δ*lysZ* (**Fig. 5B**). Complementation of the *mptB*, *manC*, and *ppmC* deletions by expressing the corresponding gene from an ectopic locus restored the observed Cog Δ*lysZ* plaque formation defect (**Fig. S3B**). Inactivation of MptA had an intermediate effect on Cog Δ*lysZ* plaque formation, while loss of MptC had little or no effect on Cog Δ*lysZ* plaque formation, respectively (**Fig. 5B**). Another use of DPM in the envelope is protein mannosylation by the mannosyltransferase Pmt^24^ (**Fig. 5A**). Deletion of *pmt* had no effect on the plaquing ability of Cog Δ*lysZ* phage (**Fig. 5B**). Notably, unlike deletion of *ppm*C or other genes required for DPM formation that both restored Cog Δ*lysZ* plaque formation and conferred resistance to LysZ toxicity, inactivation of MptB only affected Cog Δ*lysZ* plaque formation. Cells lacking MptB remained sensitive to ^Cog^*lysZ* expression (**Fig. 5C**). We therefore conclude that the function of LysZ is to overcome what we infer to be an envelope stabilizing activity of LM/LAMs with the initial phases of mannan polymerization being sufficient to prevent lysis in the absence of LysZ. By contrast, the toxic activity of LysZ only required the presence of DPM, not its downstream products, suggesting that the lysis promoting activity of LysZ may be triggered by its association with the lipid-linked mannose precursor.

### MptB inactivation likely results in depletion of the decaprenyl lipid carrier

Loss of LM/LAM production via deletion of *mptA* in *M. smegmatis* was recently shown to cause a growth and cell shape defect when cells are cultured on LB agar but not standard 7H10 medium^25^. This growth defect could be suppressed by increasing the osmolarity of the LB medium with the addition of sorbitol, indicating that the load-bearing, turgor-resistance function of the cell envelope is compromised upon MptA inactivation. We similarly observed that *Cglu* cells lacking MptB, the enzyme upstream of MptA for LM/LAM biogenesis, had shape defects and a severe growth defect on LB medium that is suppressed by sorbitol addition (**Fig. 5D**). Notably, deletion of *ppmC*, which blocks DPM formation, suppressed the growth and shape defects caused by the inactivation of MptB. Based on these results, we conclude that the severity of the Δ*mtpB* phenotype in *Cglu* is likely mediated by the sequestration of the decaprenyl lipid carrier in the DPM pool, causing a reduction in the lipid-carrier dependent process of peptidoglycan synthesis that affects cell growth and shape (see Discussion).

### Identification of the likely DPM flippase for LM/LAM biogenesis

Cgp_1254 is a predicted to be a member of the GtrA-like lipid flippase family which has been implicated in transporting polyprenyl-linked sugars across the cytoplasmic membrane^26^. Additionally, inactivation of *cgp_1254* results in a phenotypic fingerprint that is similar to that caused by *ppmC* defects^27^. Thus, Cgp_1254, which we have renamed *dpmF* is an excellent candidate for the as yet unidentified DPM flippase needed for LM/LAM biogenesis. To test this possibility, we took advantage of the Cog Δ*lysZ* mutant as a probe for the production of LM/LAM. Similar to other mutations that negatively impact LM/LAM synthesis, deletion of *dpmF* suppressed the plating defect of the Cog Δ*lysZ* phage (**Fig. 6A**). Moreover, DpmF inactivation resulted a loss of detectable LM/LAM in cell lysates that was comparable to that observed in Δ*mptB* cells with the exception that a small amount of low molecular weight glycans were observed in the Δ*dpmF* mutant (**Fig. 6B**). Accordingly, cells deleted for *dpmF* displayed an intermediate sensitivity to ^Cog^LysZ overproduction (**Fig. S4**). Cells lacking DpmF were resistant to low levels of ^Cog^*lysZ* induction, but unlike the Δ*ppmC* mutant, they were sensitive to higher levels of ^Cog^*lysZ* expression. We therefore conclude that DpmF is likely to be the dedicated flippase that transports DPM across the cytoplasmic membrane for LM/LAM biogenesis. Furthermore, in the absence of DpmF it appears that a low level of DPM transport is mediated by an alternative flippase, enabling the production of a small amount of short LM glycans and rendering cells sensitive to high levels of ^Cog^LysZ.

**Figure 6.**
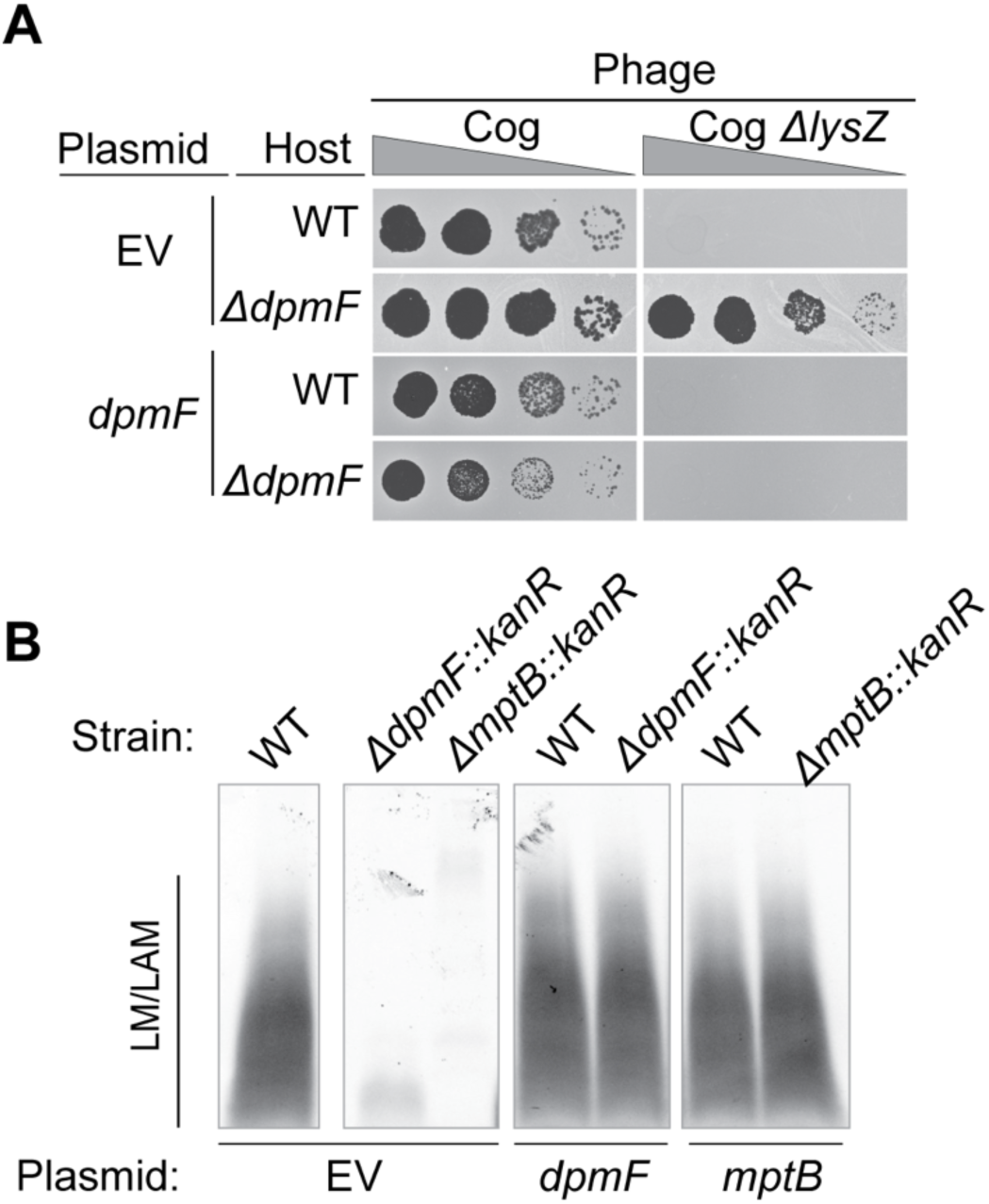
Evidence that *dpmF* encodes the DPM flippase. **A**. Tenfold dilutions of the indicated phage spotted on lawns of hosts containing the indicated expression plasmid. Expression was induced with IPTG (10 µM) and theophylline (1 mM) in cultures for approximately 6 hours before lawns were prepared and induction was maintained in the soft agar. **B**. LM/LAM was harvested from overnight cultures of the indicated strains and analyzed by SDS-PAGE and ProQ Emerald staining. Overnight cultures were induced with IPTG and theophylline as in **A**.

## DISCUSSION

Phages that infect bacteria within the Mycobacteriales order require specialized lytic functions to rupture the complex cell envelope of their hosts for the release of progeny virions. Mycobacteriophages supplement the core holin-endolysin system that destroys the peptidoglycan cell wall with LysB esterases that disrupt the mycomembrane by cleaving linkages connecting mycolic acids to the arabinogalactan layer^9,10^. The LysB requirement for efficient host cell lysis indicates that the mycomembrane of mycobacteria serves as a significant barrier to phage progeny dispersal. We were therefore surprised to find that corynephages lack identifiable *lysB*-like genes despite having hosts that also build a mycomembrane. This *lysB* deficiency prompted us to search for other corynephage genes that might encode functions targeting the mycomembrane to induce cell lysis. The *lysZ* gene emerged as a good candidate due to its broad distribution within the lysis cassettes of corynephages and other phages that infect members of the Mycobacteriales order. Indeed, deletion of *lysZ* resulted in a plaque formation and lysis defect for two corynephages Cog and CL31 that infect our model host *Cglu*. Notably, this defect was corrected by plasmid-based expression of *lysZ*-like genes from diverse phages that infect a range of different hosts, indicating that these genes encode a conserved activity required for efficient host cell lysis.

Genetic analysis of LysZ activity indicated that it is not required to disrupt the mycomembrane as we initially suspected. This result coupled with the lack of *lysB*-like genes in the lysis cassettes of corynephages suggests that the mycomembrane of corynebacteria may not be as formidable a barrier to phage release as the corresponding membrane of mycobacteria, which contains much longer mycolates that are covalently linked to the arabinogalactan layer to a greater degree than in the corynebacterial membrane^28,29^. Rather than targeting the mycomembrane, our results indicate that LysZ proteins are required to overcome a barrier created by LM/LAM lipoglycans. Cells of *Cglu* were rapidly lysed by wildtype phages. However, Cog phage lacking LysZ was not capable of efficiently lysing cells with LM/LAMs, which instead of lysing were converted to misshapen yet osmotically stable protoplast-like cells by the mutant phage. This phenotype is reminiscent of the lysis defect caused by spanin inactivation in phages that infect Gram-negative bacteria^6^. Infections with such phages yield spheroplasts stabilized by the outer membrane, a result that hinted at a stress-bearing role for the outer membrane layer of the envelope that has since been well established^7^. By analogy with the spanins, our results with LysZ and the wide distribution of *lysZ*-like genes among corynephages and mycobacteriophages suggest that LM/LAMs play a similar mechanical role in stabilizing the envelope of *Cglu* and other members of the Mycobacteriales order (**Fig. 7**).

**Figure 7.**
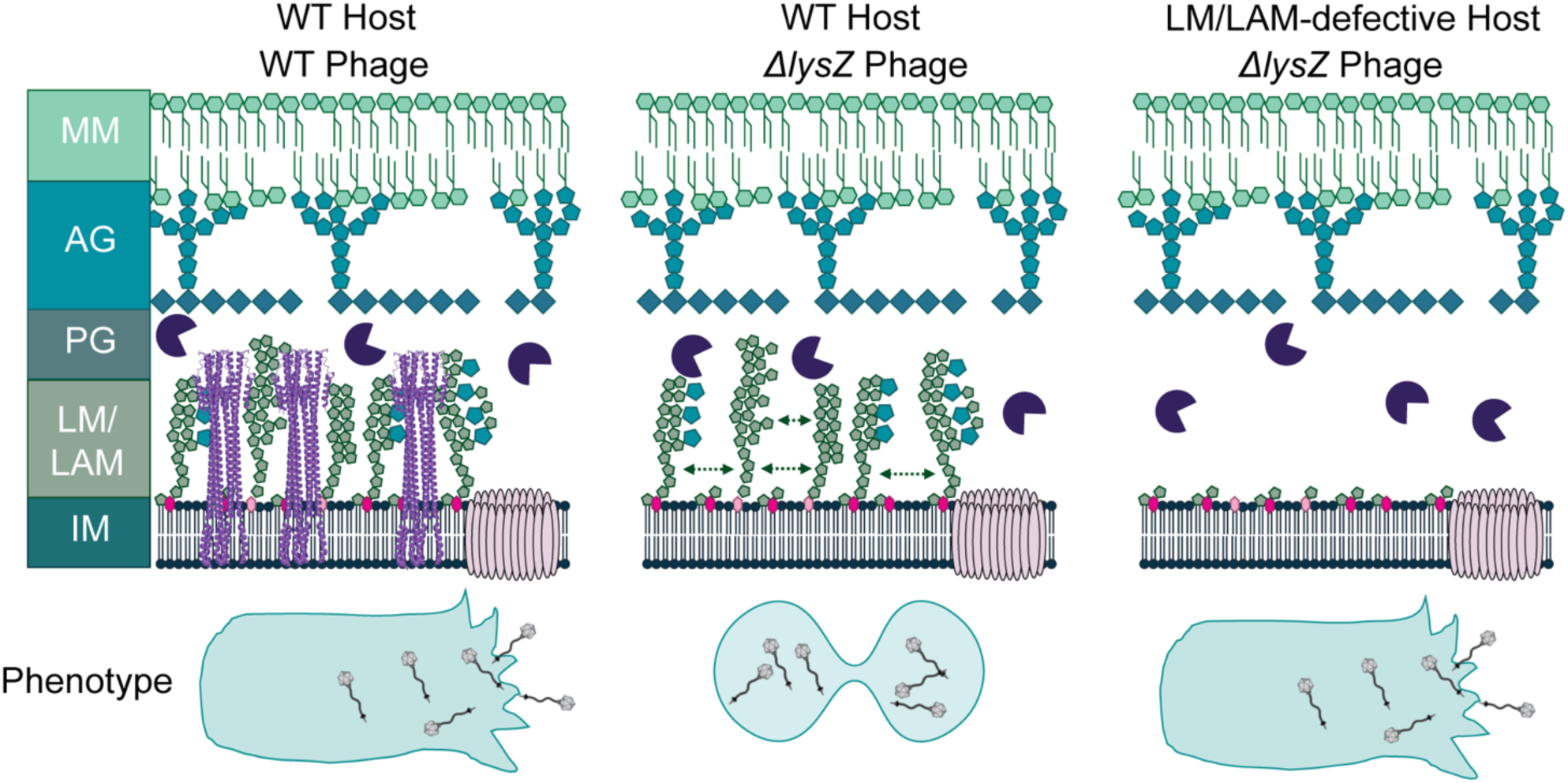
Model for LysZ function. **Left.** LysZ functions to disrupt interactions between LM/LAM polymers to assist host cell lysis following holin triggering and PG degradation by the endolysin. **Middle.** Infection of wild-type *Cglu* with a phage lacking LysZ results in lysis failure due to the stabilizing properties of LM/LAM lipoglycans. **Right.** Infections of cells lacking LM/Lam do not require LysZ to induce host cell lysis.

A recent study of LM/LAM lipoglycans in *M. smegmatis* found that cells lacking the MptA enzyme that elongates LM/LAM polymers display morphological and growth defects, suggesting a role for these molecules in cell shape and division^25^. Blocking a similarly late stage in LM/LAM synthesis in *Cglu* by inactivating the MptB enzyme also results in cell shape and division defects. However, we found that inactivating the enzyme that forms the lipid-linked LM/LAM precursor DPM suppressed this phenotype. This result indicates that the shape and division problems of Δ*mptB* mutants are not caused by the loss of LM/LAM per se but instead are the result of secondary effects stemming from the failure to complete LM/LAM synthesis. This phenomenon is commonly observed in bacterial mutants defective for the synthesis of a surface glycan. For example, defects in the later steps of teichoic acid synthesis in Firmicutes or in enterobacterial common antigen synthesis in *E. coli* cause growth and shape defects that are suppressed by inactivating the enzyme that forms the first lipid-linked intermediate in the respective pathway^30–32^. The problems caused by late-stage blocks in these pathways are thought to arise due to the shared use of the polyprenyl lipid carrier by surface glycan synthesis pathways and the pathway for peptidoglycan cell wall biogenesis. When a glycan synthesis pathway is blocked after the committed step, lipid carrier-linked intermediates of the pathway accumulate, reducing the availability of the carrier. The limited carrier supply then indirectly inhibits/reduces peptidoglycan synthesis to cause the observed growth and morphological defects. Our results with the Δ*mptB* mutants of *Cglu* suggest that the growth and shape defects induced by blocking the late stages of LM/LAM synthesis also stem from indirect effects on peptidoglycan synthesis. We suspect that the indirect inhibition of peptidoglycan synthesis also contributes to the phenotypes observed for MptA inactivation in *M. smegmatis*, but further experiments are required to test this hypothesis.

Blocking the early stages of LM/LAM synthesis by deleting *ppmC* in *Cglu* results in a growth defect. Although indirect effects are difficult to rule out, this result suggests an important role for LM/LAM polymers for cell growth that is independent of the potential problems with peptidoglycan synthesis induced by late-stage blocks in the LM/LAM synthesis pathway. Why LM/LAM polymers are important for growth is not clear, but the observation that they present a barrier to phage induced lysis suggests that their function may involve the mechanical stabilization of the cell envelope. How these polymers might stabilize the envelope remains unclear, but we envision that networks of hydrogen bonds between adjacent polymers could form a stress-bearing shell around the membrane. Thus, LM/LAM interactions may function analogously to the lateral packing of LPS molecules in the outer membrane of Gram-negative bacteria, the stress-bearing properties of which are dramatically enhanced by the long glycan chains of the O-antigen^7,33^. Bridging interactions between lipid anchored polymers related to LM/LAMs like lipoteichoic acids may also provide structural properties to the envelope of Gram-positive Firmicutes that phages of these bacteria must also disrupt to promote efficient lysis.

How LysZ overcomes the barrier to lysis presented by the LM/LAM lipoglycans remains unknown. One possibility is that the long helical structure predicted for LysZ interferes with the lateral interactions between LM/LAM polymers that are normally involved in stabilizing the membrane against osmotic rupture. Notably, the toxicity of LysZ overproduction requires DPM production. Thus, another attractive possibility is that LysZ functions similarly to the lantibiotic nisin, which forms pores in the membrane after it binds the lipid-linked PG precursor lipid II^34^. Accordingly, LysZ may similarly form a membrane disrupting pore structure following its association with DPM. Further studies of LysZ function and of phage lysis systems in general promises to reveal new insights into the biology of bacterial cell envelopes and how they can be disrupted for potential biotechnological and therapeutic applications.

## ACKNOWLEDGEMENTS

The authors would like to acknowledge all the members of the Bernhardt and Rudner labs for thoughtful discussion and generous advice. Particularly, we would like to thank Anastacia Parks for creating and sharing the *pmt* and *manA* mutants and Eric Snow for sharing and trouble-shooting LM/LAM analysis protocols. We would like to thank Dr. Graham Hatfull and the SEA-PHAGES program for sharing stocks of the phages Fury and Corndog. We also would like to thank Paula Montero Llopis and her team at the Microscopy Resources on the North Quad (MicRoN) core facility at HMS for the microscopy services. This work was supported by Investigator funds from the Howard Hughes Medical Institute, and ACM was supported in part by the Life Sciences Research Foundation where she was a Simons Fellow.

## AUTHOR CONTRIBUTIONS

A.C.M. and T.G.B. designed research; A.C.M. and E.W.L. performed research; A.C.M. and T.G.B. analyzed data; A.C.M. and T.G.B. wrote the paper.

## METHODS

### Media, strains, plasmids, and phages

*Cglu* was grown in LB supplemented with 0.4% glucose (LBG) or Brain Heart Infusion supplemented with 9.1% sorbitol (BHIS) at 30°C with aeration. Kanamycin (kan), chloramphenicol (chlor) and apramycin (apra) were added to the media as needed at concentrations of 15 μg/mL, 3 ug/mL and 12.5 μg/mL respectively. Theophylline was used at a concentration of 1 mM and IPTG was used at a concentration of 0.1 mM for induction of gene expression unless otherwise indicated. *E. coli* was grown in LB at 37°C or 30°C with aeration with 25 μg/mL kan or 50 μg/mL apra as needed. All strains, phages plasmids, and primers used are listed in Supplemental Tables S2-S5.

### Preparation of *Cglu* competent cells and electroporation

Saturated cultures of *Cglu* strains were diluted 1:100 into BHI+supplements (9.1% sorbitol and 2.5% glycine) and grown for 16 hours at 18°C. Cells were harvested by centrifugation at 3000xg for 10 minutes at 4°C. The cell pellet was then washed 3 times with cold 10% glycerol and resuspended in 10% glycerol to an approximate OD600 of 20. DNA was added to 100 μL aliquots of the electrocompetent cells on ice before being electroporated at 1.7 kV. Once electroporated, cells were immediately flushed with fresh BHIS and heat-shocked for 6 minutes at 46°C. Cells were then outgrown at 30°C for 1 hour and plated on antibiotic-containing plates.

### Plasmid and recombinant DNA construction

Details on plasmid assembly can be found in Supplementary Methods. Assembled *Cglu* plasmids were transformed into *E. coli* or electroporated directly into *Cglu*. Linear *Cglu* recombination cassettes were generated with 50 to 500 base pairs of homology upstream and downstream of the desired deletion. These cassettes were assembled either by direct amplification of a kanamycin cassette from plasmid pSEC01 or pEWL74 using primers with 50bp of homology to the chromosome and 20bp to the cassette or through amplification of the homologous regions and assembly with the kanamycin cassette using overlapping extension (SOE) PCR.

To establish a stable *^Cog^lysZ* expression plasmid, a pTGR5^35^ derivative was created with a theophylline-inducible translational *riboE1* riboswitch^36^ replacing the ribosome biding site of pTGR5 to make pACM40. ^Cog^*lysZ* was cloned into this vector via isothermal assembly (ITA) and transformed into E. coli. Upon transformation into *Cglu*, individual colonies were picked and patched in the presence and absence of inducer to confirm the maintenance of *^Cog^lysZ* toxicity. Plasmid from an isolate displaying a high degree of inducer sensitivity was then sequenced and found to have an A>G point mutation between the P*tac* and *riboE1* (pACM387). This plasmid, with expression control referred to as P*tac*\*riboE1 was then further used for expression studies throughout this work.

### Strain construction

#### Allelic exchange

Clean deletions were generated through the use of the pCRD206 temperature sensitive plasmid^37^, wherein 500-1kb fragments were amplified upstream and downstream of the desired gene. These fragments were assembled into pCRD206 via ITA. Plasmids were then electroporated into *Cglu* and grown at 25°C. Successful transformants were then purified at 30°C to select for integration. Integrants were then outgrown at 25°C and plated on 10% sucrose at 30°C to select for double-crossover events. The presence of the desired deletion allele was then validated through diagnostic PCR.

#### Recombineering

*Cglu* strains were engineered through recombineering as described by Hart et al.^38^ using a single-stranded annealing protein (SSAP) and cognate single-stranded binding protein (SSB) from a from *Troponema socranskii* prophage^39^. Briefly, the recombineering plasmid (pEWL54 or pEWL103) was electroporated into MB001. For pEWL54, cells were grown at 30°C in BHIS chlor. An overnight culture of the pEWL54 recombineering strain was then back diluted 1:50 into BHI+supplements (9.1% sorbitol and 2.5% glycine) and grown at 30°C with aeration. After ∼3 hours when the cells were mid-log, the SSAP/SSB pair were induced with 1mM PTG and grow was continued with aeration for 2 more hours. These cells were then harvested as described above. A total of 1μg of DNA was added to each 100 μL aliquot of electrocompetent cells and electroporated as above. Cells were plated on BHIS kan and grown at 30°C. Colony PCR was used to determine successful recombination events, and colonies were patched in the presence and absence of chlor to monitor loss of the recombineering plasmid.

For pEWL103, which is temperature-sensitive for replication, cells were grown at 25°C in BHIS apra. An overnight culture of the pEWL103 strain was then back diluted 1:50 and grown at 25°C with aeration in BHI+supplements. After ∼4 hours when the cells were mid-logarithmic phase, the SSAP/SSB pair was then induced with 1mM theophylline and 1 mM IPTG and growth was continued with aeration for 4 more hours. These cells were then harvested as described above and 1 μg DNA template was added to each 100 μL aliquot of electrocompetent cells. Cells were electroporated as above and outgrown for 2 hours at 25°C. Cells were then plated on BHIS kan at 25°C. Colony PCR was used to determine successful recombination events, and colonies were then grown at 30°C to cure the recombineering plasmid. To make an unmarked deletion, strains were recombineered with a kanR cassette flanked by *loxP* sites as above. Once confirmed, such mutants were electroporated with pCre and grown for two days at 25°C. Colonies were then purified and grown at 30°C overnight to cure the plasmid. Individual colonies were then picked and patched onto media with and without kan or apra to identify transformants that lost both the resistance cassette and the pCre plasmid. Colony PCR was used to determine successful recombination events.

### Bacterial viability assays

The optical density of overnight bacterial cultures was normalized to OD600=1 and serially diluted 10-fold. A volume of 5 μl of each dilution was spotted onto a plate and allowed to dry. Plates were incubated at 30°C for one or two nights before imaging.

### Bacterial growth kinetics

Overnight cultures of *Cglu* were diluted 1:100 into LBG apra and grown to midlog. Cultures were normalized to an OD600=0.15 and added to a 96-well plate with 0.1 mM IPTG and 1 mM theophylline in biological triplicates and technical duplicates. Optical density was measured every 10 minutes with shaking at 30°C over the course of 16 hours in a Tecan Nano.

### Bacterial colony forming unit (CFU) measurement

Overnight cultures of *Cglu* were diluted 1:100 into LBG apra and grown to midlog. Cultures were normalized to an OD600=0.4 and serially tenfold diluted. A volume of 100 μL of each dilution was plated for single colonies in technical duplicates and incubated overnight at 30°C. The normalized cultures were induced with 0.1 mM IPTG and 1 mM theophylline overnight. In the morning, the OD600 for each culture was recorded and serially diluted. A volume of 100 μL of each dilution was plated for single colonies in technical duplicates and incubated overnight at 30°C. Fold chance in CFU was calculated by dividing the CFU post-induction by the CFU pre-induction. Experiments were performed in biological triplicates.

### Phage propagation and genetic manipulation

#### *Cglu* phages

For isolation and enumeration of phage stocks, *Cglu* hosts were grown to mid-log (OD600 = 0.3-0.5) in LBG supplemented with 10mM CaCl2 and 100 μL cells were mixed with 10 μL of phage diluted in SM buffer and allowed to adsorb for 10 minutes. The mixture was then added to LBG 0.5% agar supplemented with 10 mM CaCl2, spread on normal agar, and allowed to solidify. Solid plates were incubated overnight at 30°C. Confluent lysis plates were used to grow high titer phage stocks. These plates were overlayed with SM buffer and gently rocked overnight at 4°C. Overlays were then harvested and filter sterilized before being concentrated in a centrifugal column (Amicon 100 kDa).

Mutations in *lysZ* were achieved through modification of a Cas12 system^40^ (see Plasmid Construction). First, WT phage was plated onto midlog *Cglu* grown in LBG 10mM CaCl2 containing a multi-copy plasmid encoding the desired mutations with ∼200bp of homology on either side. Ten plaques from this host were picked and pooled. Next, successfully recombined phages were selected on *Cglu* containing a CRISPR-Cas12 system with a spacer against the *lysZ* allele of interest. For this step, the cells were grown in LBG 10 mM CaCl2 with 3 μg/mL chlor and the CRISPR-Cas12 system was induced with 1 mM IPTG and 1 mM theophylline. To enable selection of Cog Δ*lysZ* phages, the CRISPR-Cas plasmid pACM351 was electroporated into *Cglu* Δ*cgp_1254* and used as a Δ*lysZ* permissive host. Individual plaques were picked and validated by colony PCR and Sanger sequencing. Desired plaques were then repurified 2x on the CRISPR host and stored in SM buffer.

### Phage plaque formation assays

Phage spot assays with hosts encoding toxic lysis genes were performed by concentrating 2 mL of a *Cglu* overnight with apra into 500 μL. Then, 200 μL concentrated *Cglu* culture was added to 4 mL molten top LBG 0.5% agar supplemented with 10 mM CaCl2, 12.5 μg/mL apra, 0.1 mM IPTG and with or without 1 mM theophylline as indicated in the figure legends, vortexed, applied over a solid LBG 1.5% agar plate, and allowed to dry under a flame for 10 minutes. Phage spot assays for mannose mutants were performed by concentrating 4 mL of a mid-log *Cglu* culture into 500 μL. Then, 200 μL concentrated *Cglu* culture was added to 4 mL molten top LBG 0.5% agar supplemented with 10 mM CaCl2, vortexed, applied over a solid LBG 1.5% agar plate, and allowed to dry under a flame for 10 minutes. Phage spot assays for complementation of mannose mutants were performed by subculturing an overnight culture into fresh media with apra, 1mM theophylline, and 0.01 mM or 0.1 mM IPTG as indicated in the figure legend, growing to mid-log. Then, 4 mL of these cultures were concentrated into 500 μL and 200 μL *Cglu* culture was added to 4 mL molten top LBG 0.5% agar supplemented with 10 mM CaCl2, apra, 1 mM theophylline, and 0.01 mM or 0.1 mM IPTG, vortexed, applied over a solid LBG 1.5% agar plate, and allowed to dry under a flame for 10 minutes. Phage were diluted tenfold, and spots of 3 μL of each dilution were applied to the solidified top agar and allowed to dry under flame. Dried plates were then incubated overnight at 30 C. Phage plaque assays were performed by subculturing overnight cultures into fresh LBG with apra and 10 mM CaCl2 and outgrown to midlog. Then, 100 μL of cells and 10 μL of phage dilutions were mixed and allowed 10 minutes to adsorb. This mixture was then plated with LBG 0.25% or 0.5% top agar as labeled with 10 mM CaCl2, apra, 1 mM theophylline, and 0.1 mM IPTG. The plates were then allowed to solidify before being incubated overnight at 30°C.

### Phage lysis kinetics

Overnight *Cglu* cultures were subcultured 1:100 into LBG supplemented with 10 mM CaCl2 and apra and grown to midlog. Cultures were normalized to an OD600=0.25 and added into a 96 well plate. Phage was added at an MOI of 2.5 and cells were induced upon infection with addition of 0.1 mM IPTG and 1 mM theophylline. The plate was incubated in the Tecan Nano at 30°C with shaking, and the OD600 was recorded every 5 minutes. Samples were run in biological triplicate and technical duplicate.

### Enumeration of phages produced during infection

Overnight *Cglu* cultures were subcultured 1:100 into LBG supplemented with 10 mM CaCl2 and apra and grown to midlog. Cultures were normalized to an OD600=0.3 and phage was added at an MOI of 2.5 while inducing expression of cloned plasmid genes with 0.1 mM IPTG and 1 mM theophylline. Infected cultures were then incubated for 2 hours at 30°C. At 2 hours post-infection, 100 μl of each sample was collected and centrifuged for 3 minutes at 10,000xg to remove the unlysed cells. The supernatant was then tenfold diluted in SM buffer and tittered via dribble plating. Briefly, 10 mL of LBG 0.5% agar were mixed with 10 mM CaCl2, 1 mM theophylline, 0.1 mM IPTG and a midlog *lysZ* complement host. This mixture was poured into a petri dish, allowed to solidify under flame, and 10 μl of phage dilutions were then dribbled across the top of the solidified plate and dried under flame. Plates were then incubated overnight at 30°C and plaques were enumerated the next day.

### Phage-infected *Cglu* microscopy

Overnight *Cglu* cultures were diluted 1:100 into LBG supplemented with 10 mM CaCl2 and grown to midlog. Phage was added at an MOI of 10 and allowed 30 minutes for adsorption at 30°C with aeration. Infected cells were then spotted onto an 2% LBG agarose pad supplemented with 10 mM CaCl2 and imaged on a Nikon Ti-E inverted widefield microscope with a motorized stage, a perfect focus system, and a 1.45 NA Plan Apo ×100 Ph3 DM objective lens with Cargille Type 37 immersion oil. Images were acquired with an Andor Zyla 4.2 Plus sCMOS camera (65 nm pixel size) using Nikon Elements acquisition software (v5.10) and rendered for publication with Fiji.

### Suppressor isolation and sequencing

WT *Cglu* containing the *^Cog^lysZ* expression plasmid were grown overnight in the presence of antibiotics. Cultures were serially tenfold diluted and 100 μl of the dilutions were plated onto BHIS agar plates supplemented with 0.1 mM IPTG, 1 mM theophylline, and apra. Plates were incubated for two days at 30°C. Individual colonies were patched onto BHI plates with or without 0.008% SDS. Colonies that failed to grow on SDS but grew successfully on BHIS were then grown up in LBG supplemented with 10 mM CaCl2 over-day along with a WT control. Then, 200 μl of these cultures were used to seed lawns for a phage spot assay as described above and incubated overnight. *Cglu* suppressors that gained susceptibility to Cog Δ*lysZ* plaque formation were further purified and their genomic DNA (gDNA) was isolated. Briefly, overnight cultures were harvested and incubated with 50 mM EDTA and 1 mg/ml lysozyme at 37°C for 30 minutes to remove the cell wall. Following lysozyme treatment, the gDNA was extracted using the Promega Wizard DNA Extraction Kit and sent to the Microbial Genome Sequencing Center in Pittsburgh, PA (now named SeqCenter). Mutational analysis was performed using the breseq pipeline^41^ and Geneious Prime Workbench, and the results were validated using Sanger sequencing.

### Lipomannan and lipoarabinomannan analysis

Overnight cultures of *Cglu* were grown in BHIS supplemented with apra, 1mM theo and 0.01mM IPTG. Samples were normalized to an OD600∼5 and 1mL was pelleted and resuspend in PBS and Laemmli buffer. Samples were then boiled at 100°C for 25 minutes and run on a 4-20% TGX gel. LM and LAM were then stained using the ProQ Emeraled 300 Lipopolysaccharide Gel Stain Kit (Thermo Scientific) using standard protocols.

### Bioinformatics

Putative *lysZ*-like alleles were identified through analysis of lysis cassettes of Actinophages available on PhagesDB^42^. Representative phages from different clusters were arbitrarily chosen and visualized with Pharmerator^43^. Genes proximal to the predicted endolysin(s) as well as genes at the end of the structural modules were screened for predicted TM domains with DeepTMHMM^44^. Functional predictions were made with HHPred and Foldseek^45^ based on Alphafold2^46^ generated models. Graphs were created with GraphPad Prism and cartoons and figures were prepared in Adobe Illustrator.

## Supplemental Material

**Figure S1.**
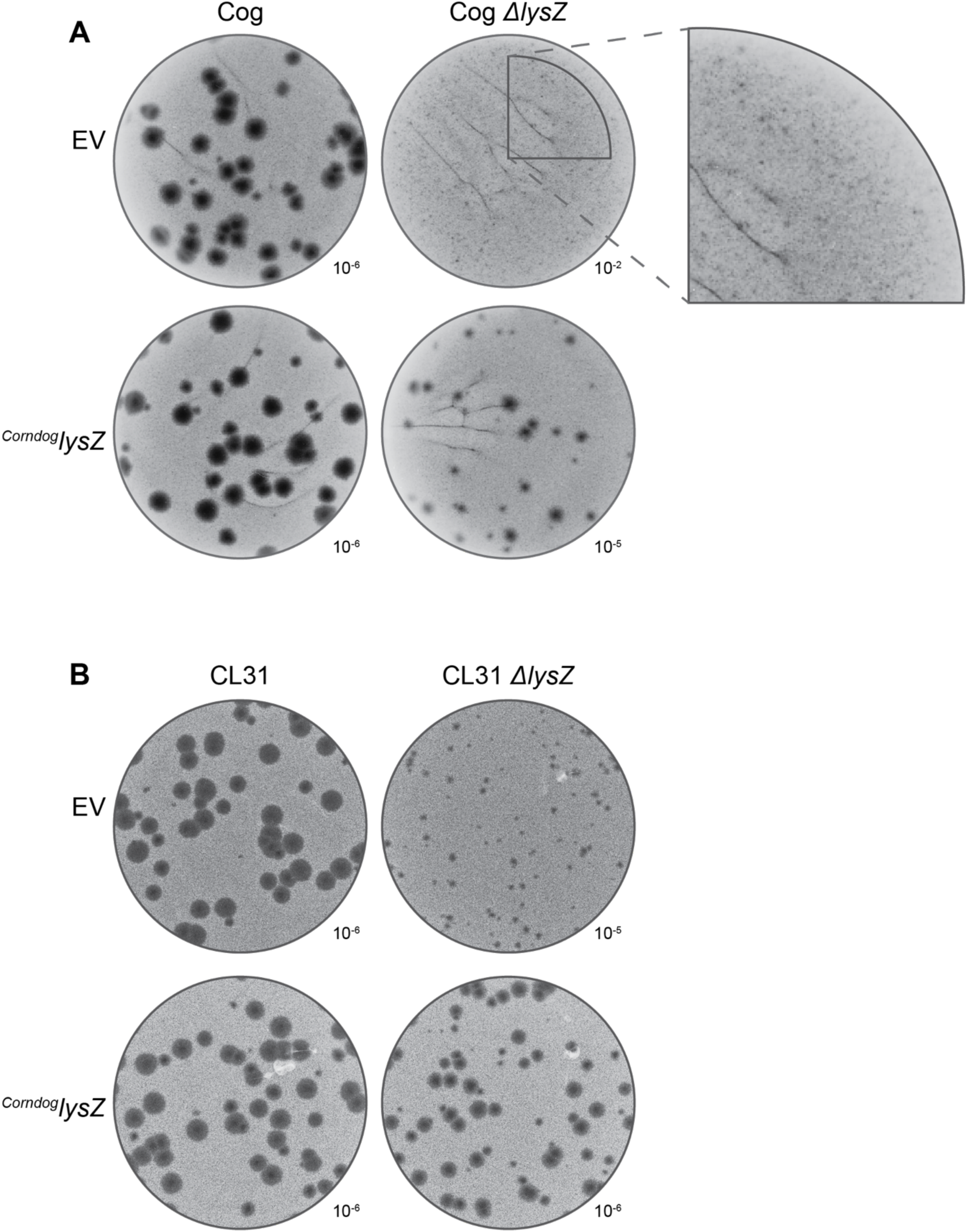
Phages lacking LysZ have defects in plaque formation. **A,** Listed dilutions of the indicated phages infecting the indicated *Cglu* expression strain in 0.25% top agar. Inset shows zoomed in view of Cog ΔlysZ plaques. Genes were induced with IPTG (100 µM) and theophylline (1 mM) in the soft agar. **B,** Listed dilutions of the indicated phages infecting the indicated *Cglu* expression strain in 0.5% top agar. Genes were induced with IPTG (100 µM) and theophylline (1 mM) in the soft agar.

**Figure S2.**
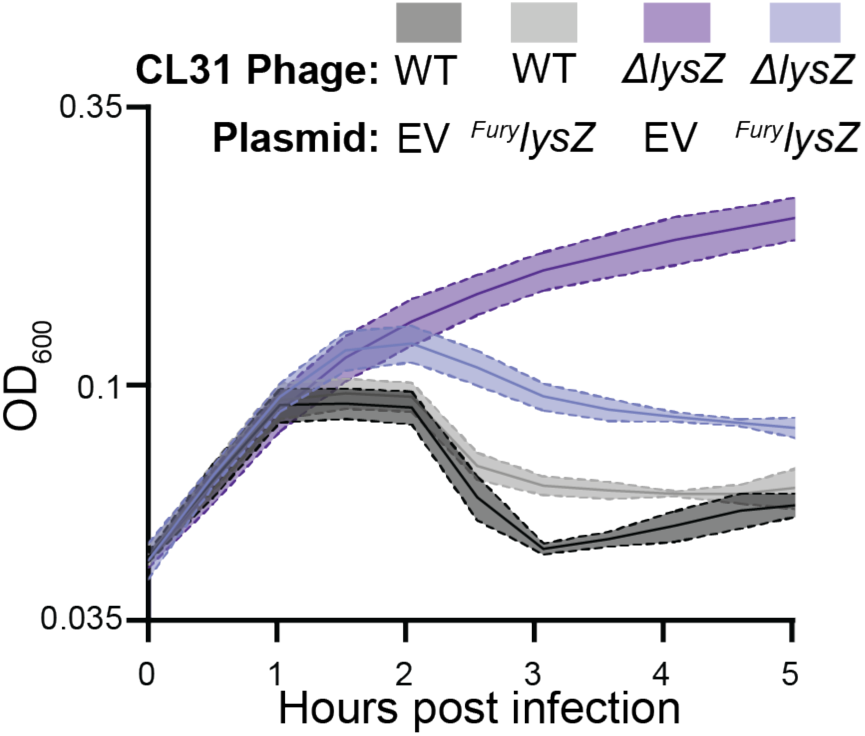
Lysis kinetics of CL31 phage. Cells of *Cglu* harboring the indicated expression plasmid were infected with the indicated phage at an MOI of 2.5. Expression of the indicated genes was induced with IPTG (100 µM) and theophylline (1 mM) at the time of infection. Growth of the infected culture was followed by monitoring OD600. Shaded regions indicate standard deviation, and solid lines indicate the mean from three independent experiments.

**Figure S3.**
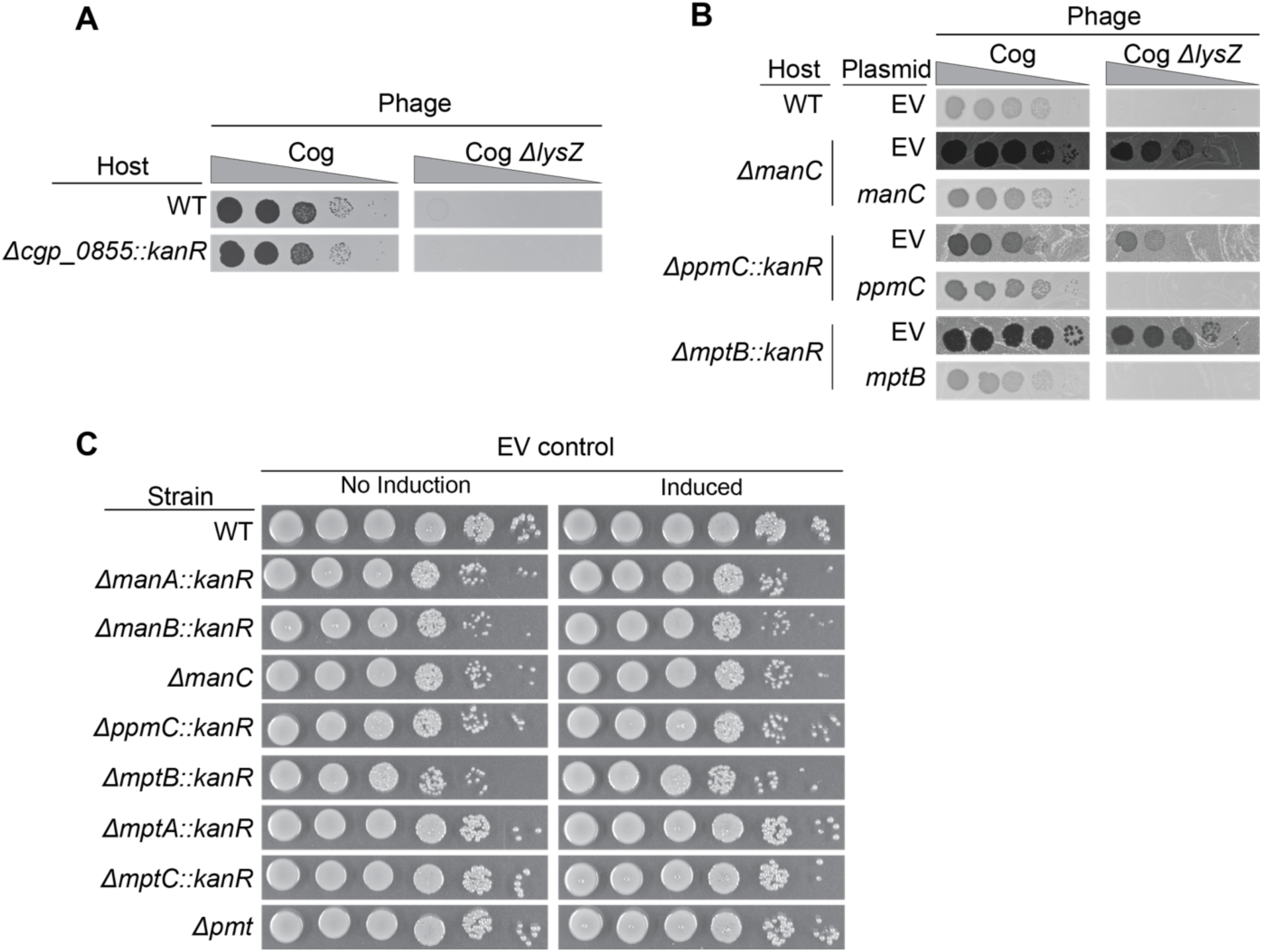
Complementation and other controls. **A**. Tenfold dilutions of the indicated phage variants spotted on lawns of the indicated mutant *Cglu* hosts. **B.** Tenfold dilutions of the indicated phage variants spotted on lawns of the indicated mutant *Cglu* hosts harboring the indicated expression plasmids. The indicated genes were under control of P*tac* and riboE1*. Induction was performed in liquid cultures for approximately 6 hours before plating with IPTG (100 µM) and theophylline (1 mM) maintained in the top agar. **C**. Tenfold dilutions of the indicated *Cglu* strains harboring the empty expression vector were grown on agar with no inducer or with IPTG (100 µM) and theophylline (1 mM).

**Figure S4.**
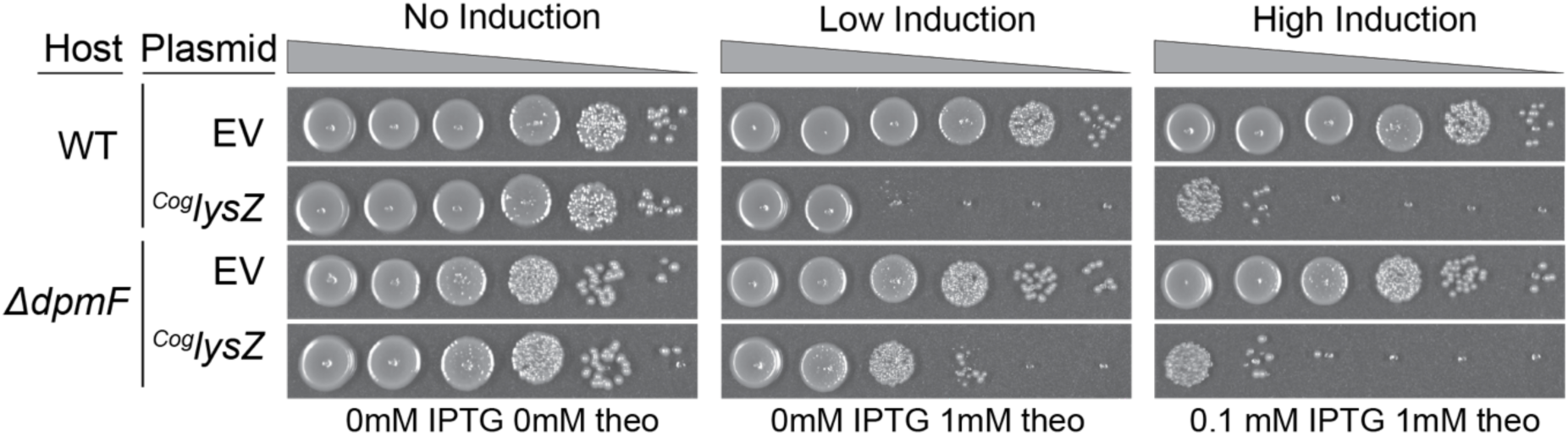
Intermediate sensitivity of *ΔdpmF* cells to *LysZ*. Tenfold dilutions of the indicated *Cglu* strains harboring the empty or ^Cog^*lysZ* expression vector were grown on agar with no inducer or with low (IPTG = 10 µM), theophylline = 1 mM) or high (IPTG = 100 µM), theophylline = 1 mM) concentrations of inducer.

## SUPPLEMENTARY MOVIES

**Movie 1. Lysis of *Cglu* infected with WT Cog.** Cells were infected with Cog at an MOI = 10. After 30 minutes for adsorption, infected cells were spotted onto an agarose pad for observation. Images were captured every 15 seconds.

**Movie 2. Lysis of *Cglu* infected with Cog *ΔlysZ* – site 1.** Cells were infected with Cog *ΔlysZ* infected and imaged as in Movie 1 except that images were captured every 1 minute.

**Movie 3. Lysis of *Cglu* infected with Cog *ΔlysZ* - site 2.** Cells were infected with Cog *ΔlysZ* infected and imaged as in Movie 1 except that images were captured every 1 minute.

## SUPPLEMENTARY TABLES

**Table S1.** *LysZ* suppressors.

| Isolate | Position | Mutation | Annotation | Gene | Description |
| --- | --- | --- | --- | --- | --- |
| Suppressor 1 | 788,382 | transposition |  | *cgp_0855* | transposon insertion in *cgp_0855*, upstream of *manA* |
| Suppressor 2 | 786,820 | A→C | D248A<br>(GAT→GCT) | *pmmA* | phosphomannomutase |
| Suppressor 2 | 980,716 | C→G | A187P<br>(GCA→CCA) | *cgp_1053* | putative transcriptional regulator, TetR-family |
| Suppressor 2 | 1,668,571 | G→T | P280Q<br>(CCA→CAA) | *gap* | glyceraldehyde-3-phosphate dehydrogenase (GAPDH) |
| Suppressor 3 | 1,547,187 | G→A | Q190*<br>(CAA→TAA) | *ppmC* | polyprenol-phosphate-mannose synthase domain 1 |

**Table S2.** Strains used in this study.

| Strain | Name | Genotype | Ref. |
| --- | --- | --- | --- |
| DH5a (Ipir) |  | *F<sup>-</sup> hsdR17 deoR recA1 endA1 phoA supE44 thi-1 gyrA96 relA1 Δ(lacZYA-argF)U169 φ80dlacZΔM15 \*\*\*\*add pir* | Gibco BRL |
| MB001 | H#0060 | MB001 (ATCC 13032 ΔCGP1 (cg1507-gp1524) ΔCGP2 (cg1746-1752) ΔCGP3 (cg1890-cg2071)) | (1) |
| *ΔahfA* | H#2262 | MB001 *ΔahfA* (*cgp_0475*) | (2) |
| *ΔotsA* | H#1666 | MB001 *ΔotsA* (*cgp_2907*) | (2) |
| Suppressor 1 | H#5306 | MB001 pTGR5_SNP_riboE-Cog_gp25 apraR *cgp_0855* transposon insertion | This paper |
| Suppressor 2 | H#5308 | MB001 pTGR5_SNP_riboE-Cog_gp25 apraR; *pmmA* asp248ala; *cgp_1053* ala187pro; *gap* pro280gln | This paper |
| Suppressor 3 | H#5307 | MB001 pTGR5_SNP_riboE-Cog_gp25 apraR; *ppmC* Gln190* | This paper |
| *Δcgp_0855::kanR* | H#5365 | MB001 *Δcgp0855::kanR lox* | This paper |
| *ΔmanA::kanR* | H#2559 | MB001 *ΔmanA::kanR lox* | This paper |
| *ΔmanB::kanR* | H#3681 | MB001 *ΔpmmA::kanR lox* | This paper |
| *ΔmanC* | H#3678 | MB001 *ΔmanC* | This paper |
| *ΔppmC::kanR* | H#3679 | MB001 *ΔppmC::kanR* | This paper |
| *ΔdpmF::kanR* | H#3680 | MB001 *ΔmanA::kanR* | This paper |
| *ΔdpmF* | H#5406 | MB001 *ΔmanA::lox* | This paper |
| *ΔmptB::kanR* | H#2439 | MB001 *ΔmptB::kanR* | This paper |
| *ΔmptB* | H#5405 | MB001 *ΔmptB::lox* | This paper |
| *ΔmptB ΔppmC::kanR* | H#5413 | MB001 *ΔmptB::lox ΔppmC::kanR* | This paper |
| *ΔmptB ΔdpmF::kanR* | H#5102 | MB001 *ΔmptB::lox ΔmanA::kanR* | This paper |
| *ΔmptA::kanR* | H#2438 | MB001 *ΔmptA::kanR* | This paper |
| *ΔmptC::kanR* | H#2440 | MB001 *ΔmptC::kanR* | This paper |
| *Δpmt* | H#2060 | MB001 *Δpmt* (*cgp_1014*) | This paper |

**Table S3.** Phages used in this study.

| Isolate | Strain no | Genotype/notes | Ref. |
| --- | --- | --- | --- |
| Cog | ACMΦ008 |  | (2) |
| Cog Δ*lysZ* | ACMΦ159 | deletion of *lysZ* (*gp25*) | This paper |
| CL31 | ACMΦ002 |  | (2) |
| CL31 Δ*lysZ* | ACMΦ095 | deletion of *lysZ* (*gp22*) | This paper |

**Table S4.** Plasmids used in this study.

| Construct name in paper | Plasmid | Genotype | Ref. |
| --- | --- | --- | --- |
| pCRE | pEWL89 | pCRD206(ApraR)::P<sub>SOD</sub>-cre | (3) |
| recombineering 1 | pEWL054 | pTGR5(CamR)::P<sub>tac</sub>-SSAP/SSB | (3) |
| recombineering 2 | pEWL103 | pCRD206(ApraR)::P<sub>tac</sub> riboE1-SSAP/SSB | (3) |
| pJYS3 | pJYS3 |  | (4) |
| pFSC | pFSC |  | (5) |
| P<sub>tac</sub>-GFP | pTGR5 | KanR, P<sub>tac</sub>-eGFP, pGA1 mini replicon | (6) |
| pCRD206::pmt | pARP1 | KanR, pCRD206 derivative containing an insert covering upstream and downstream of *cgp_1014* (pmt) | This paper |
| pCRD206::ΔmanC | pACM407 | KanR, pCRD206 derivative containing an insert covering upstream and downstream of *cgp_0422* (manC) | This paper |
| P<sub>SOD</sub>_cas12 | pEWL001 | KanR, pK-PIM derivative encoding P<sub>SOD</sub>_riboE1-Cas12 | This paper |
| P<sub>SOD</sub>_Cas12_P<sub>tac</sub>_anti-CL31<sup>lysZ</sup> spacer | pACM363 | CmR, pTGR5 derivative encoding the CRISPR-Cas12 system and a spacer to target CL31 *lysZ* | This paper |
| P<sub>SOD</sub>_Cas12_P<sub>tac</sub>_anti-Cog<sup>lysZ</sup> spacer | pACM351 | CmR, pTGR5 derivative encoding the CRISPR-Cas12 system and a spacer to target Cog *lysZ* | This paper |
| P<sub>SOD</sub>_Cas12_P<sub>tac</sub>_CRISPR-array | pEWL018 | CmR, pTGR5 derivative encoding P<sub>SOD</sub>_Cas12 and P<sub>tac</sub>_CRISPR array with BsaI digest sites for spacer editing | This paper |
| P<sub>SOD</sub>_dCas12_P<sub>tac</sub>_CRISPR-array | pEWL013 | CmR, pTGR5 derivative encoding P<sub>SOD</sub>_dCas12 and P<sub>tac</sub>_CRISPR array with BsaI digest sites for spacer editing | This paper |
| P<sub>tac</sub>_riboE<sub>manA</sub> | pACM964 | ApraR, pTGR5 derivative encoding P<sub>tac</sub>_riboE1-manA | This paper |
| P<sub>tac</sub>_riboE<sub>CL31<sup>lysZ</sup></sub> | pACM490 | ApraR, pTGR5 derivative encoding P<sub>tac</sub>_riboE1-CL31<sup>lysZ</sup> (gp22) | This paper |
| P<sub>tac</sub>_riboE<sub>Cog<sup>lysZ</sup></sub> | pACM385 | KanR, pTGR5 derivative encoding P<sub>tac</sub>_riboE1-Cog<sup>lysZ</sup> (gp25) | This paper |
| P<sub>tac</sub>_riboE<sub>Cog<sup>lysZ</sup></sub> | pACM413 | ApraR, pTGR5 derivative encoding P<sub>tac</sub>_riboE1-Cog<sup>lysZ</sup> (gp25) | This paper |
| P<sub>tac</sub>_riboE<sub>Corndog<sup>lysZ</sup></sub> | pACM583 | ApraR, pTGR5 derivative encoding P<sub>tac</sub>_riboE1-Corndog<sup>lysZ</sup> (gp75) | This paper |
| P<sub>tac</sub>_riboE<sub>D29<sup>lysB</sup></sub> | pACM519 | ApraR, pTGR5 derivative encoding P<sub>tac</sub>_riboE1-D29<sup>lysB</sup> (gp12) | This paper |
| P<sub>tac</sub>_riboE<sub>EV</sub> | pACM414 | ApraR, pTGR5 derivative encoding P<sub>tac</sub>_riboE1 empty vector | This paper |
| P<sub>tac</sub>_riboE<sub>Fury<sup>lysZ</sup></sub> | pACM489 | ApraR, pTGR5 derivative encoding P<sub>tac</sub>_riboE1-Fury<sup>lysZ</sup> (gp59) | This paper |
| P<sub>tac</sub>_riboE<sub>manC</sub> | pACM488 | ApraR, pTGR5 derivative encoding P<sub>tac</sub>_riboE1-manC | This paper |
| P<sub>tac</sub>_riboE<sub>mptB</sub> | pACM514 | ApraR, pTGR5 derivative encoding P<sub>tac</sub>_riboE1-mptB | This paper |
| P<sub>tac</sub>_riboE<sub>ppmC</sub> | pACM491 | ApraR, pTGR5 derivative encoding P<sub>tac</sub>_riboE1-ppmC | This paper |
| P<sub>tac</sub>_riboE<sub>Smarties<sup>lysZ</sup></sub> | pACM594 | ApraR, pTGR5 derivative encoding P<sub>tac</sub>_riboE1-Smarties<sup>lysZ</sup> (gp53) | This paper |
| P<sub>tac</sub>_CRISPR-array | pEWL003 | CmR, pTGR5 derivative encoding P<sub>tac</sub>-CRISPR array with BsaI digest sites for spacer editing | This paper |
| P<sub>tac</sub>_EV | pACM352 | KanR, pTGR5 derivative encoding P<sub>tac</sub>_empty vector | This paper |
| P<sub>tac</sub>_riboE<sub>Cog<sup>lysZ</sup></sub> | pACM377 | KanR, pTGR5 derivative encoding P<sub>tac</sub>_riboE1-Cog<sup>lysZ</sup> | This paper |
| P<sub>tac</sub>_riboE-GFP | pACM040 | KanR, pTGR5 derivative encoding P<sub>tac</sub>_riboE1-GFP | This paper |
| P<sub>tac</sub>-GFP | pACM246 | CmR, pTGR5 derivative encoding P<sub>tac</sub>-GFP | This paper |
| CL31 Δ<sup>lysZ</sup> editing plasmid | pACM321 | CmR, pTGR5 derivative encoding CL31 editing template to delete *lysZ* | This paper |
| Cog Δ<sup>lysZ</sup> editing plasmid | pACM376 | CmR, pTGR5 derivative encoding Cog editing template to delete *lysZ* | This paper |

\*A>G SNP between P<sub>tac</sub> and riboE1.

**Table S5.**
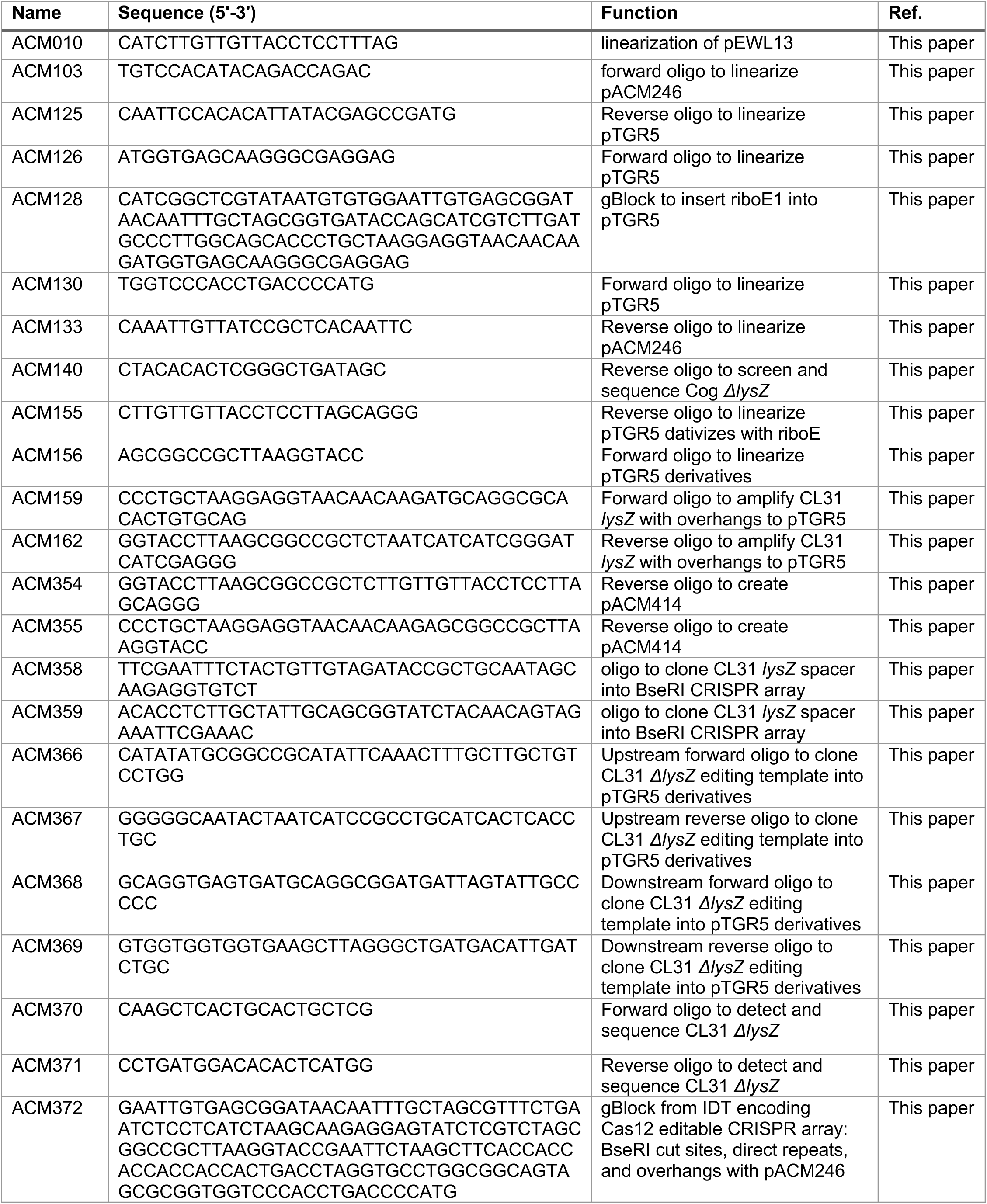
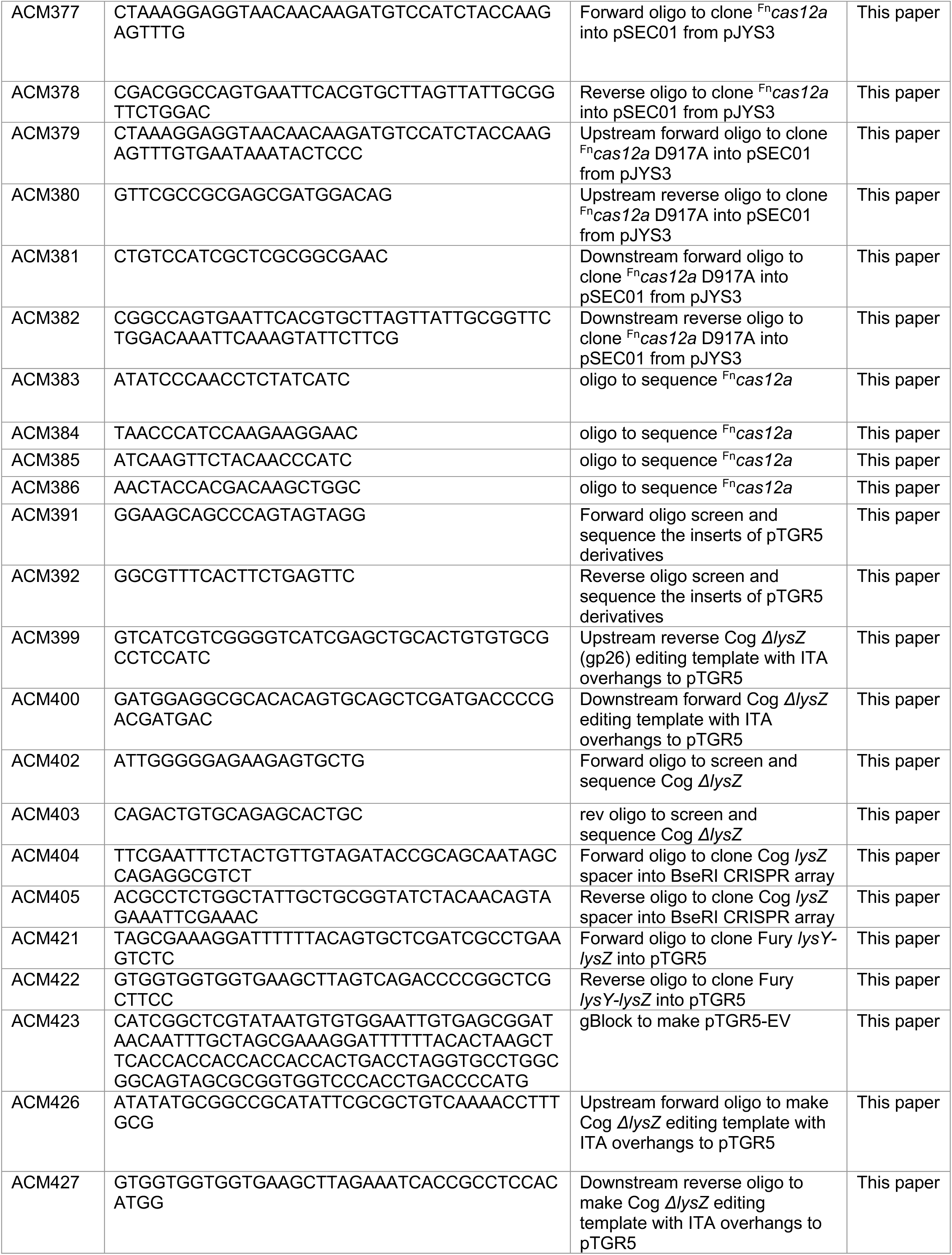
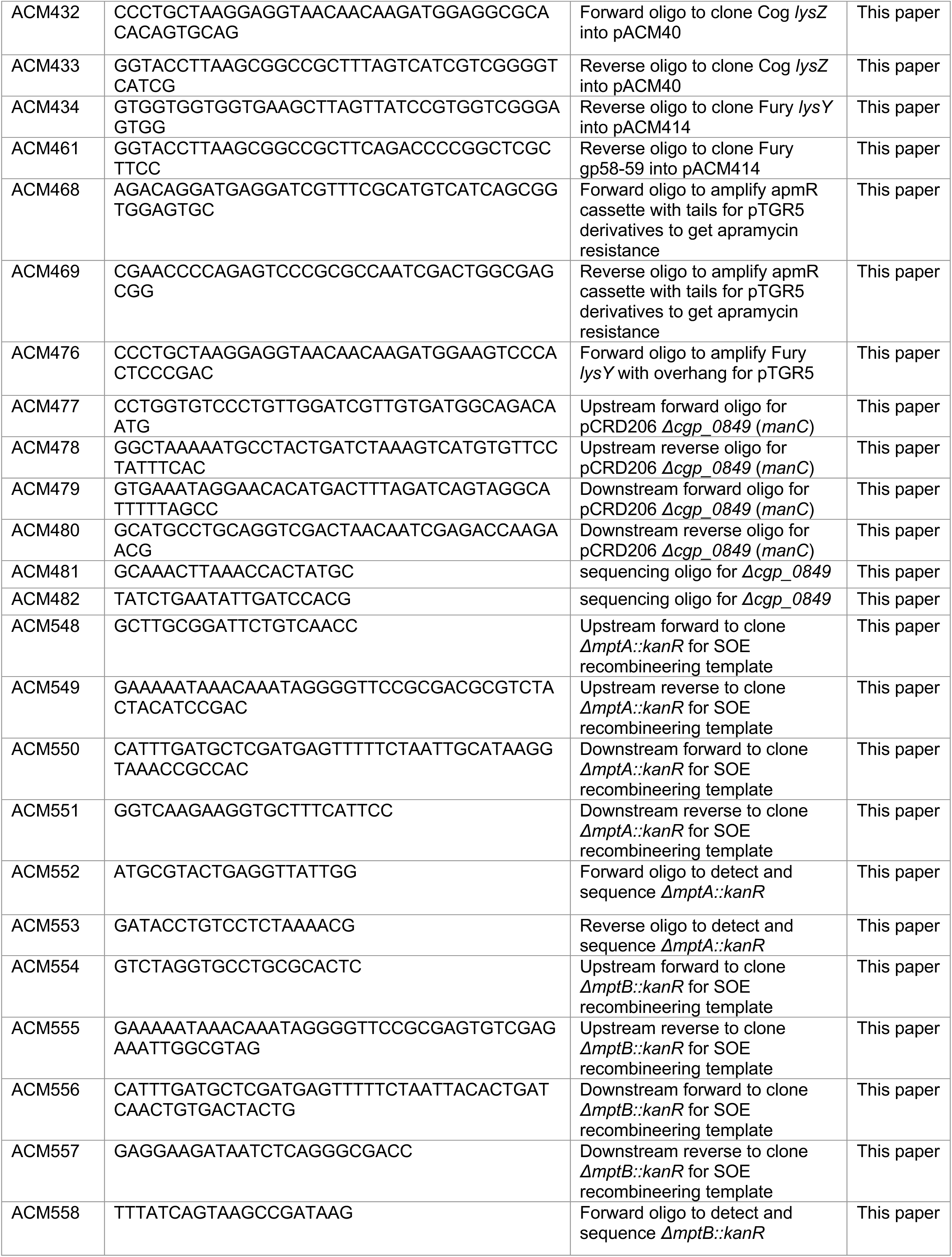
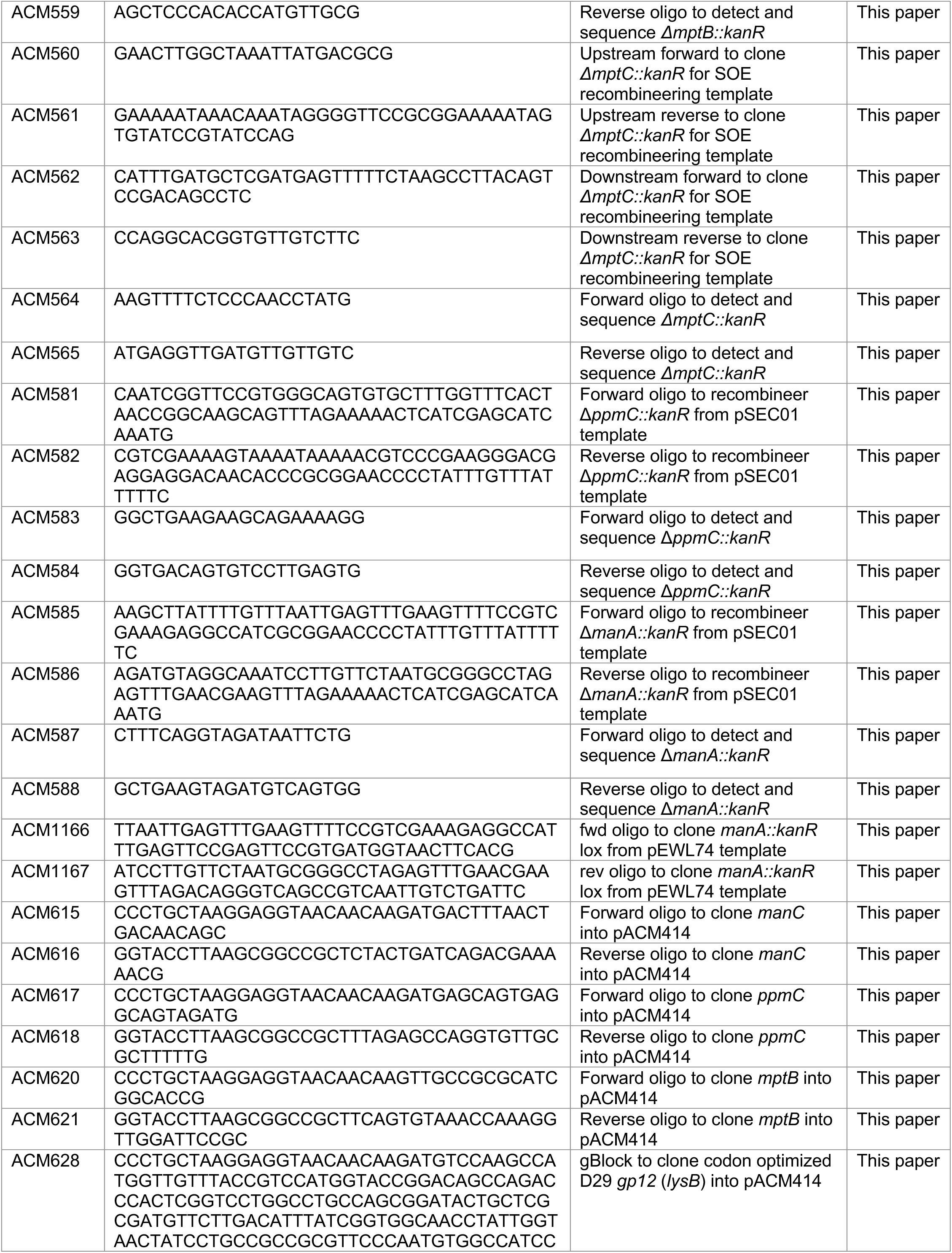
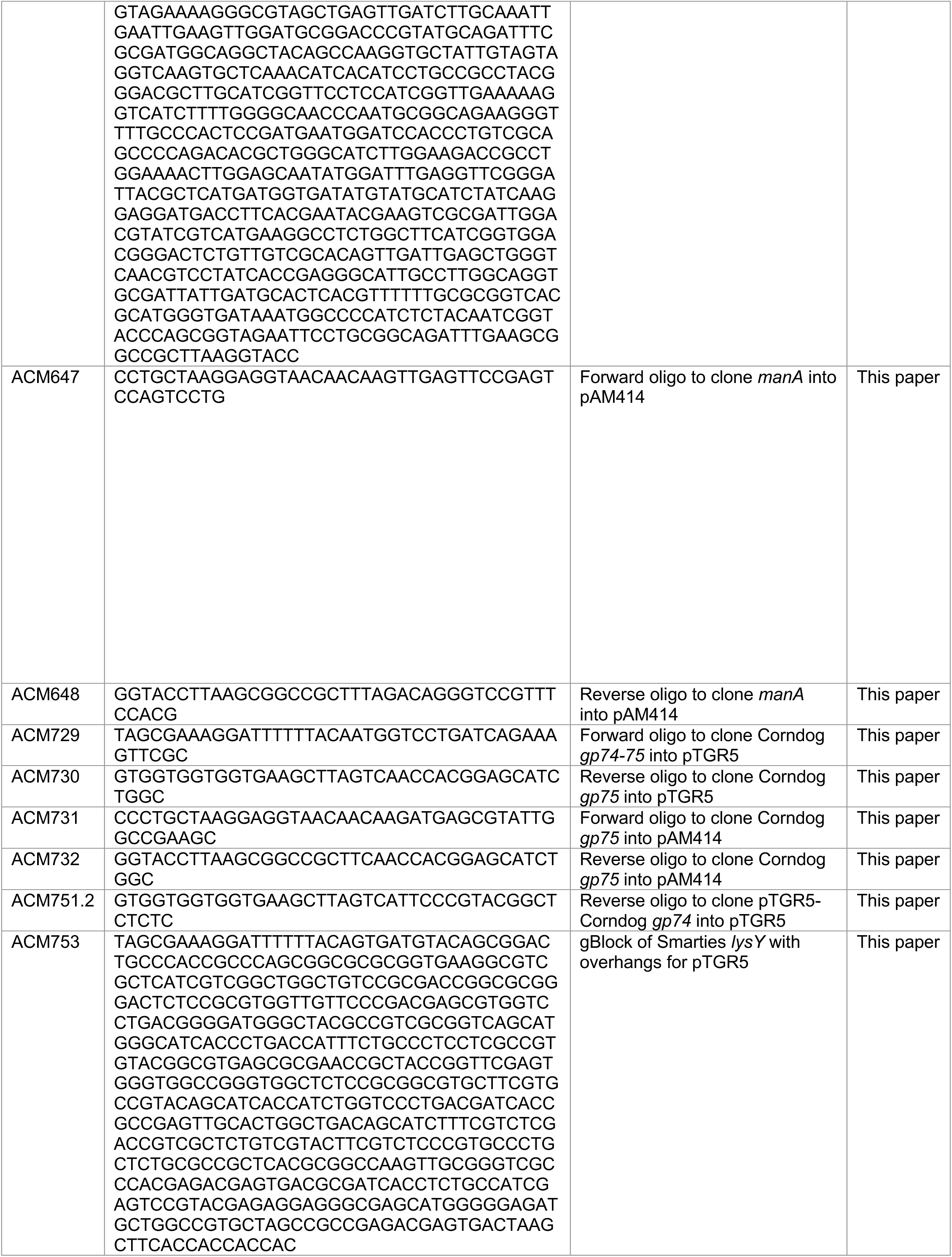
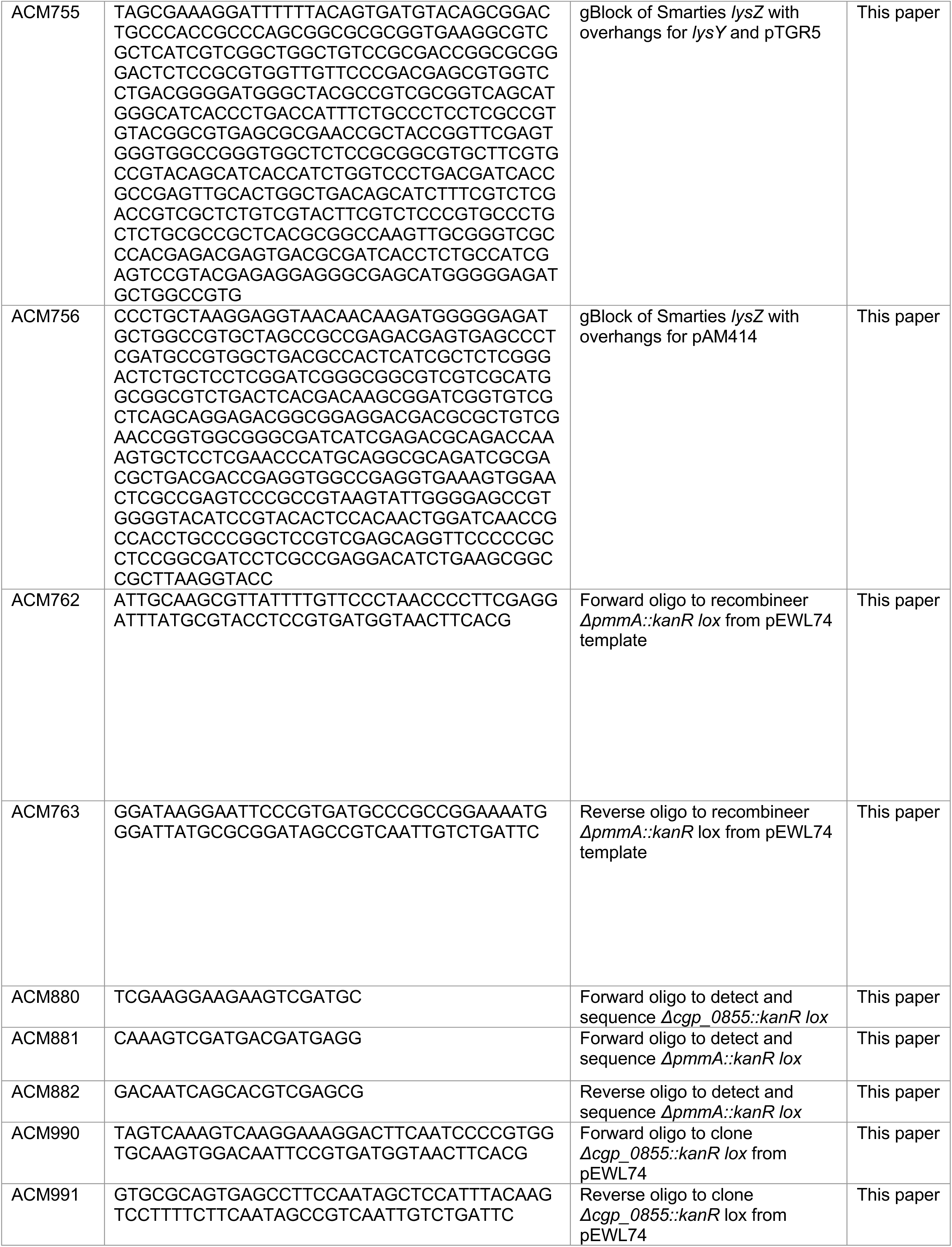
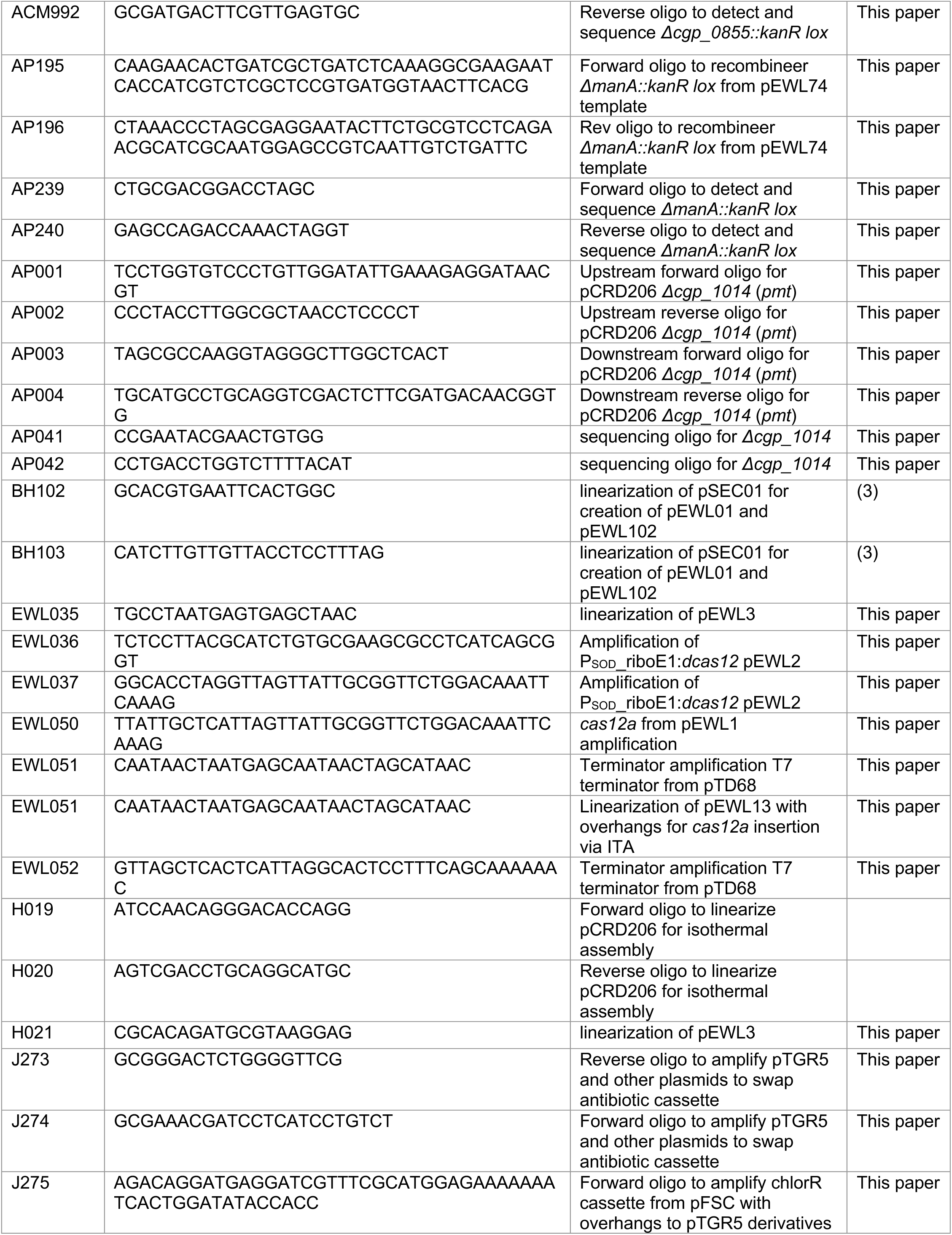
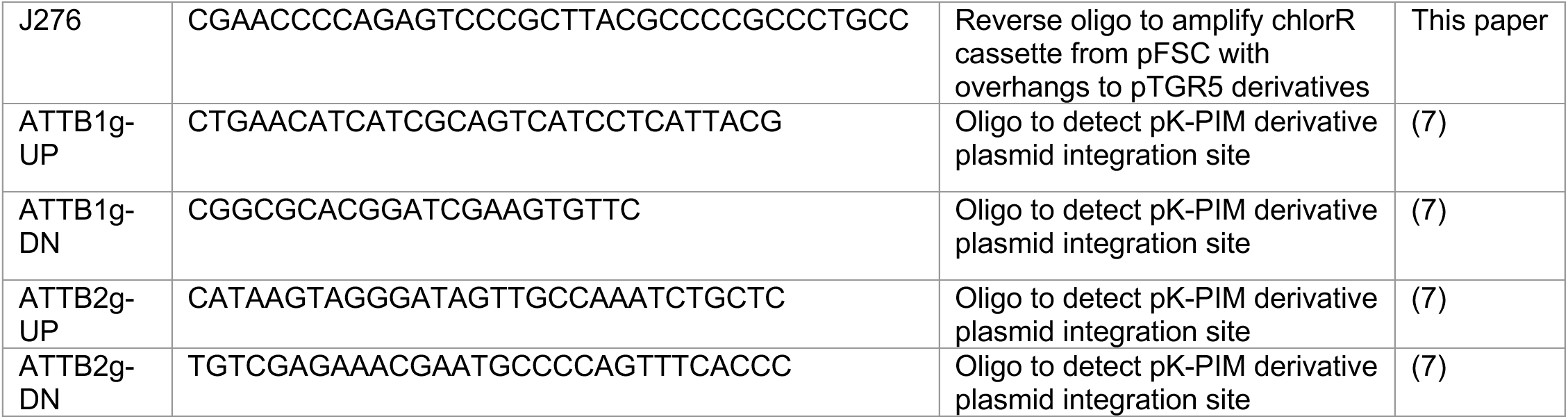
Primers used in this study.

## SUPPLEMENTAL MATERIALS AND METHODS

### Plasmid Construction

#### pCRD206::*pmt* (pARP1)

PCR amplified pCRD206 backbone (AP53-54), 750bp upstream of *cgp_1014* and first 3 amino acids (AP1-2), and 750bp downstream including last 6aa of *cgp_1014* (AP3-4), assembled via ITA.

#### pCRD206::*ΔmanC* (pACM407)

PCR amplified pCRD206 backbone (H19/H20), ∼1kb upstream and 3 amino acids (ACM477/478), and ∼1kb downstream including last 2aa (ACM479/480), assembled via ITA.

#### PSOD_*cas12* (pEWL001)

The vector pSEC1 was amplified using the primers BH102 and BH103. The Franciscella novicida *cas12* gene was amplified from pJYS3 using the primers ACM377 and ACM378 and assembled via ITA. pJYS3_ΔcrtYf was a gift from Sheng Yang (4) (Addgene plasmid # 85542 ; http://n2t.net/addgene:85542 ; RRID:Addgene_85542).

#### PSOD_*cas12*_Ptac_anti-^CL31^*lysZ*_spacer (pACM363)

Oligos ACM358/359 were annealed, PNK treated, and inserted into the pEWL018 CRISPR array via Golden Gate assembly using BseRI.

#### PSOD_ *cas12*_Ptac_anti-^Cog^*lysZ*_spacer (pACM351)

Oligos ACM404/405 were annealed, PNK treated, and inserted into the pEWL018 CRISPR array via Golden Gate assembly using BseRI.

#### PSOD_ *cas12*_Ptac_CRISPR-array (pEWL018)

The vector pEWL3 was linearized with the primers EWL35 and H21. The catalytically inactive Francisella novicida *dcas12* under control of PSOD-riboE1 was amplified from pEWL2 with the primers EWL36 and EWL37. The second insert contained a T7 terminator amplified from pTD68 with the primers EWL51 and EWL52. The vector and two inserts were assembled via ITA.

#### PSOD_d *cas12*_Ptac_CRISPR-array (pEWL013)

The vector pACM246 was linearized with the following primers: ACM103 and ACM133. The CRISPR array containing the Type IIS restriction enzyme BseRI cut sites was ordered as a gBlock (ACM372) from IDT. The vector and gblock were ligated by ITA.

#### Ptac_*riboE_*manA* (pACM964)

PCR amplified pAM414 backbone (ACM155/156) and *manA* from MB001 gDNA (ACM647/648), assembled via ITA.

#### Ptac_*riboE_^CL31^*lysZ* (pACM490)

PCR amplified pAM414 backbone (ACM155/156) and gp22 (*lysZ*) from CL31 gDNA (ACM159/162), assembled via ITA.

#### Ptac_*riboE_^Cog^*lysZ* (pACM385)

pACM377 electroporated into MB001. Only got suppressors. Colony was picked and struck on BHI +/- IPTG theophylline to ensure that construct was still inducible. Sequencing revealed an A>G SNP between Ptac and riboE (*riboE).

#### Ptac_*riboE_^Cog^*lysZ* (pACM413)

pACM385 linearized (J274/274) and apramacying resistance casette from pFSC (ACM468/469), assembled via ITA.

#### Ptac_*riboE_^Corndog^*lysZ* (pACM583)

PCR amplified pAM414 backbone (ACM155/156) and *gp75* (*lysZ*) from Corndog gDNA (ACM731/732), assembled via ITA.

#### Ptac_*riboE_^D29^*lysB* (pACM519)

PCR amplified pAM414 backbone (ACM155/156) and *lysB* from D29 codon optomized gBlock ACM628, assembled via ITA.

#### Ptac_*riboE_EV (pACM414)

pACM413 linearized (ACM354/355) with overlaps to create an empty vector, assembled via ITA.

#### Ptac_*riboE_^Fury^*lysZ* (pACM489)

PCR amplified pAM414 backbone (ACM155/156) and *gp59* (*lysZ*) from Fury gDNA (ACM476/461), assembled via ITA.

#### Ptac_*riboE_*manC* (pACM488)

PCR amplified pAM414 backbone (ACM155/156) and *manC* from MB001 gDNA (ACM615/616), assembled via ITA.

#### Ptac_*riboE_*mptB* (pACM514)

PCR amplified pAM414 backbone (ACM155/156) and *mptB* from MB001 gDNA (ACM620/621), assembled via ITA.

#### Ptac_*riboE_*ppmC* (pACM491)

PCR amplified pAM414 backbone (ACM155/156) and *ppmC* from MB001 gDNA (ACM617/618), assembled via ITA.

#### Ptac_*riboE_^Smarties^*lysZ* (pACM594)

PCR amplified pAM414 backbone (ACM155/156) and *gp53* (*lysZ*) from Smarties in gBlock ACM756, assembled via ITA.

#### Ptac_^Corndog^*lysYZ* (pACM589)

pTGR5 digested with NdeI and EcoRI and *gp74-75* (lysY/*lysZ*) amplified from Corndog gDNA with ACM729/730, assembled via ITA.

#### Ptac_^Corndog^*lysY* (pACM593)

pTGR5 digested with NdeI and EcoRI and *gp74* (lysY) amplified from Corndog gDNA with ACM729/751.2, assembled via ITA.

#### Ptac_CRISPR-array (pEWL003)

The vector pSEC1 was amplified using the primers BH102 and BH103. The Francisella novicida cas12 gene was amplified in two fragments from pJYS3 in order to introduce a D917A catalytic mutation. The first fragment utilized primers ACM379 and ACM380, while the second set of primers included ACM381 and ACM382. The vector and two inserts were assembled by ITA.

#### Ptac_EV (pACM352)

PCR amplified pTGR5 backbone (ACM125/130) and gBlock ACM423, assembled via ITA.

#### Ptac_^Fury^*lysY* (pACM379)

pTGR5 digested with NdeI and EcoRI and *gp58* (*lysY*) amplified from Fury gDNA with ACM421/434, assembled via ITA.

#### Ptac_^Fury^*lysYZ* (pACM349)

pTGR5 digested with NdeI and EcoRI and *gp58-59* (*lysY*-*lysZ*) amplified from Fury gDNA with ACM421/422, assembled via ITA.

#### Ptac_riboE-^Cog^*lysZ* (pACM377)

PCR amplified pAM040 backbone (ACM155/156) and *lysZ* (gp26) from Cog gDNA (ACM432/433), assembled via ITA.

#### Ptac_riboE1-GFP (pACM040)

PCR amplified pTGR5 backbone (ACM125/126) and gBlock ACM128 with riboE1, assembled via ITA.

#### Ptac_^Smarties^*lysY* (pACM592)

pTGR5 digested with NdeI and EcoRI and Smarties *gp52* (*lysY*) gBlock (ACM753) was assembled via ITA.

#### Ptac_^Smarties^*lysYZ* (pACM595)

pTGR5 digested with NdeI and EcoRI and gBlocks (ACM755/756) encoding *gp52* and *gp53* (*lysY-lysZ*) were assembled via ITA

#### Ptac-GFP (pACM246)

PCR amplified pTGR5 backbone (J273/274) and chloroform resistance cassette (J275/276) from pFSC (5)

#### Δ^CL31^*lysZ* editing plasmid (pACM321)

Digested pTGR5 with XbaI and EcoRI (removed Ptac and RBS), amplified ∼200bp upstream and 3 amino acids (ACM366/367) and ∼200bp downstream including last 2 amino acids (ACM368/369) of *gp22* (*lysZ*), assembled via ITA.

#### Δ^Cog^*lysZ* editing plasmid (pACM376)

Digested pTGR5 with XbaI and EcoRI (removed Ptac and RBS), amplified ∼200bp upstream and 6 amino acids (ACM426/399) and ∼200bp downstream including last 7 amino acids (ACM400/427) of Cog *lysZ* (*gp25*), assembled via ITA.

